# The genomic basis and environmental correlates of local adaptation in the Atlantic horse mackerel (*Trachurus trachurus*)

**DOI:** 10.1101/2022.04.25.489172

**Authors:** Angela P. Fuentes-Pardo, Edward D. Farrell, Mats E. Pettersson, C. Grace Sprehn, Leif Andersson

## Abstract

Understanding how populations adapt to local environments is increasingly important to prevent biodiversity loss due to climate change. Here we examined whole-genome variation of twelve Atlantic horse mackerel samples from the North Sea to North Africa, and the western Mediterranean Sea. This marine migratory benthopelagic fish is one of the most widely distributed and commercially important species in the eastern Atlantic. We found low population structure at neutral loci, but high differentiation at adaptive loci distinguishing the western Mediterranean and the North Sea populations from other Atlantic locations. Candidate genes distinctive of the Mediterranean include a green-sensitive-opsin harbouring two missense mutations that might fine-tune the spectral sensitivity to blue-green light conditions. Candidate genes characteristic of the North Sea could play a critical role in cold tolerance (energy metabolism and cell membrane structure) and increased sensitivity to odours, presumably to compensate reduced visibility in turbid waters. We also discovered a putative chromosomal inversion (9.9 Mb) that follows a climate-related latitudinal cline with a break near mid Portugal. Genome-environment association analysis indicated that seawater-dissolved oxygen concentration and temperature are likely the main environmental drivers of local adaptation. Our genomic data broadly supports the current stock divisions, but recommends revision of the western and southern stock boundaries. We developed a reduced SNP panel that genetically discriminate the North Sea and North Africa from neighbouring populations. Our study highlights the importance of life history and chromosomal inversions in adaptation with gene flow, and the complexity of evolutionary and ecological processes involved in local adaptation.

## 1. Introduction

The extent to which marine species with no evident barriers restricting gene flow show genetic differentiation and local adaptation across geographic areas is a question of considerable interest in evolutionary biology, conservation, and management (Palumbi, 1994). Several marine species exhibit large population sizes, high gene flow, and minute genetic drift, resulting in low genetic differentiation that has been difficult to resolve with neutral genetic markers (Hauser & Carvalho, 2008a).

Owing to advances in high-throughput sequencing, recent genomic studies screening thousands to millions of genetic markers across the genome have revealed population structure and selection signatures in species previously assumed to be panmictic [e.g., Atlantic herring (Han et al., 2020)] or lowly structured [e.g., Atlantic cod (Barth et al., 2017), Atlantic halibut (Kess et al., 2021)]. Population structure in marine fish has been characterized by shifts in allele frequencies at many small effect loci or fewer large effect loci (Gagnaire & Gaggiotti, 2016) and in chromosomal rearrangements (Akopyan et al., 2022; Han et al., 2020; Matschiner et al., 2022). Moreover, genomic divergence has been linked to ecological diversity, for example, in migratory behavior (Kirubakaran et al., 2016), seasonal reproduction (Lamichhaney et al., 2017; Petrou et al., 2021), or along environmental gradients (Han et al., 2020; Stanley et al., 2018). Therefore, a thorough examination of genomic variation, including neutral and adaptive loci, can help identify distinct biological units and genetic variants associated with adaptation to local environments, knowledge of great interest in conservation and management, especially in the face of climate change.

Fish stock identification is an important prerequisite for fisheries management (Cadrin & Secor, 2009), however many exploited stocks have traditionally been defined according to geographical and political features or regions rather than on a biological basis. As more information becomes available, it is evident that the temporal and spatial distributions of most fisheries resources are not aligned to these artificial divisions (Kerr et al., 2017) and that biological populations are more dynamic and complex (Reiss, Hoarau, Dickey-Collas, & Wolff, 2009; Stephenson, 2002). Therefore, it is critical to identify the underlying population structure and use this information to identify the appropriate level at which to define assessment and management units. It is also important to be able to assign individuals in mixed surveys and commercial catches to the population or assessment unit to which they belong (Casey, Jardim, & Martinsohn, 2016; Hintzen et al., 2015).

The Atlantic horse mackerel (*Trachurus trachurus* Linnaeus, 1758) is a marine benthopelagic shoaling species widely distributed in the east Atlantic, from Norway to southern Africa (FAO Major Fishing areas 27, 34 and 47) and in the Mediterranean Sea (FAO Major Fishing area 37) (Froese & Pauly, 2021). It inhabits a variety of environments and different populations may be under contrasting selection pressures, making this species ideal for the study of local adaptation. Horse mackerel are generally found in continental shelf waters (100-200 m depth), but have also been observed in deeper (∼1000 m) or near-shore waters. The species undertakes annual migrations between spawning, feeding and over-wintering areas (Abaunza et al., 2003). Horse mackerel are considered to be asynchronous batch spawners with indeterminate fecundity and are not known to be faithful to their original spawning grounds (Gordo et al., 2008; Ndjaula, Hansen, Krüger-Johnsen, & Kjesbu, 2009). Eggs and larvae are pelagic and are typically either found over the continental shelf, from the surface to 100 m depth, or near the coast (Alvarez & Chifflet, 2012; Beveren, Klein, Serrão, Gonçalves, & Borges, 2016).

In the northeast Atlantic, horse mackerel are assessed and managed as three separate stocks: the Western, the North Sea, and the Southern (Figure S1), which were largely defined based on the results of the HOMSIR project (Abaunza et al., 2008). Populations inhabiting coastal waters along Morocco and Mauritania, in northwest Africa, are considered a separate group, denominated the “Saharo-Mauritanian stock”. However, these populations are less studied and monitored than those in the north. The discreteness of the stocks, as well as the location and levels of mixing between them, is unknown, which leads to uncertainty in the input data for stock assessments. Previous genetic studies have indicated low genetic differentiation between populations, and have provided inconclusive results in regards to population sub-structuring beyond the three main stock divisions identified using traditional methods (Brunel et al., 2016; Cimmaruta, Bondanelli, Ruggi, & Nascetti, 2008; Comesaña, Martínez-Areal, & Sanjuan, 2008; Farrell & Carlsson, 2018; Healey et al., 2020; Kasapidis & Magoulas, 2008; Mariani, 2012; Sala Bozano et al., 2015).

Given the elusive nature of the population structure of the Atlantic horse mackerel and its ecological and commercial importance in the east Atlantic, the aims of this study were four-fold: (i) Estimate the extent of genetic differentiation between populations based on whole-genome sequencing; (ii) Identify the genetic basis of local adaptation; (iii) Identify which ecological and evolutionary processes may have shaped patterns of population subdivision; (iv) Design a cost-effective and informative SNP panel that can be used for future population studies and genetic stock identification.

## 2. Materials and Methods

### 2.1 Sampling and DNA isolation

Samples were collected opportunistically between 2015 and 2017 through existing fishery surveys, fisheries targeted to horse mackerel, and as bycatch at 12 locations across the east Atlantic and the western Mediterranean Sea (Figure 1). We aimed to collect spawning fish to ensure that samples could provide a valid baseline (Figure S1). However, due to the opportunistic nature of sampling this was not always possible (Table S1). Maturity stages were recorded by sample collectors using a number of different maturity keys. Therefore, these were standardised to the six-point international horse mackerel maturity scale (Table S2; (ICES, 2015)). A 0.5 cm^3^ piece of tissue was excised from the dorsal musculature of each specimen and stored at 4°C in absolute ethanol. Total genomic DNA was extracted from the majority of samples by Weatherbys Scientific Ltd, Ireland from 30 mg of tissue using sbeadex™ magnetic bead-based extraction chemistry on the LGC Oktopure™ platform. The remaining samples were extracted using a CTAB or Chelex and proteinase-K based extraction protocol (Table S1). The Chelex protocol produced single-stranded DNA, whereas the other methods, double-stranded. DNA quantity was measured with a NanoDrop ND-1000 spectrophotometer.

**Figure 1.**
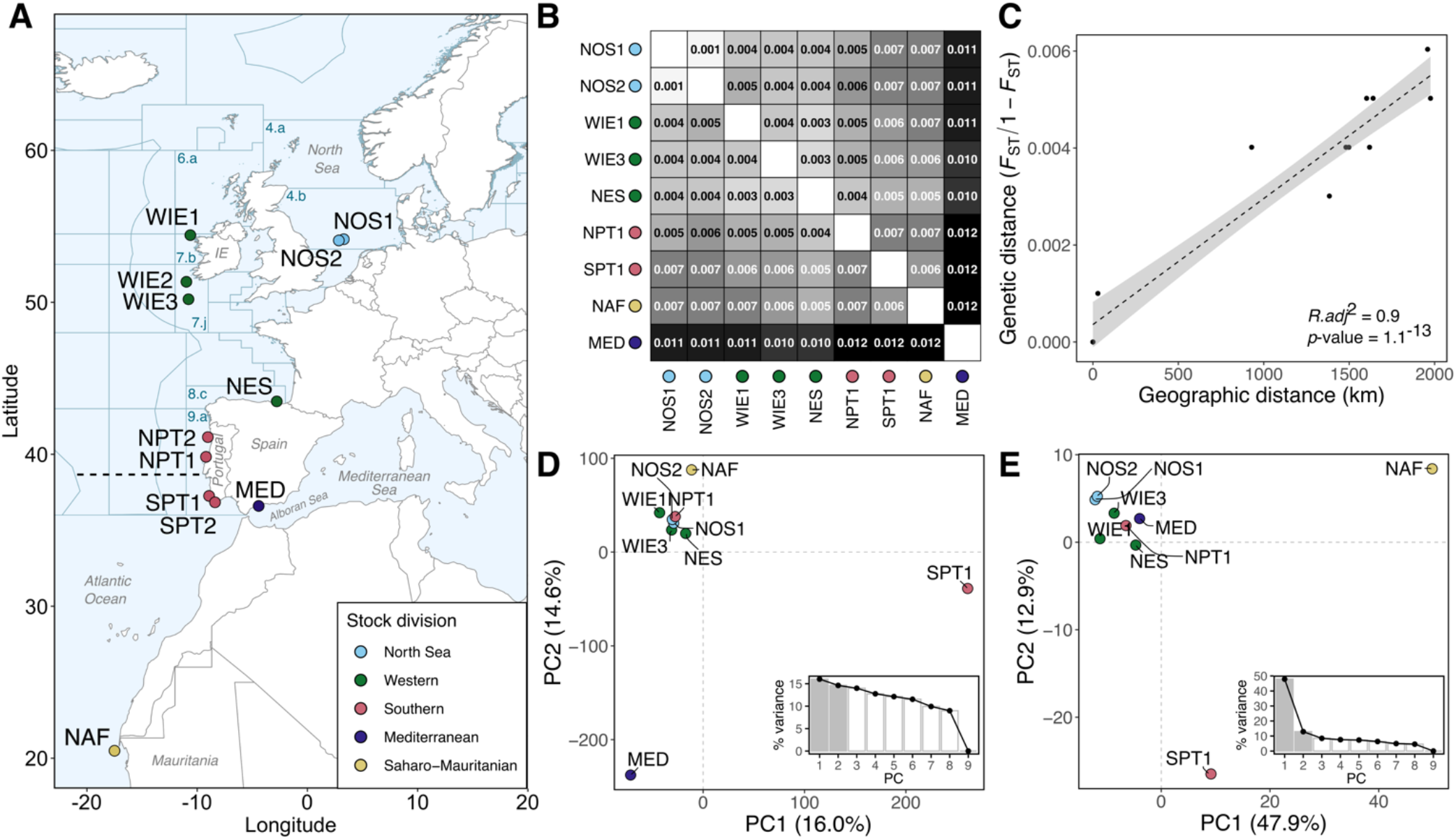
Sampling sites and population structure of the Atlantic horse mackerel. (**A**) Map depicting the 12 sampling sites in the east Atlantic Ocean. ICES fishing divisions are denoted with dark blue lines, same as their alphanumerical code. The approximate location of a biogeographical transition zone north of Lisbon, in Portugal (∼38.7-39.0°N), is denoted with a horizontal dashed line (Cunha, 2001; Santos et al., 2007). In all plots, each dot represents a sampling location and its color indicates the corresponding ICES stock division (ICES, 2005) after the HOMSIR project (Abaunza et al., 2008). (**B**) Heatmap representing pairwise pool-*F*_ST_ values based on ∼12 million SNPs. (**C**) Isolation by distance (IBD) plot illustrating the relationship between genetic distances (linearized pairwise pool-*F*_ST_) and geographic distances in km for the Atlantic populations in the “northern” group. The dashed line indicates the linear relationship between variables. (**D-E**) Principal component analysis (PCA) plot based on (**D**) undifferentiated (61,543 SNPs) and (**E**) highly differentiated (818 SNPs) markers. The first two axes are shown. Inset bar plots in each PCA plot show the percentage (%) of genetic variance explained by the first nine principal components (PC). Note that samples WIE3, NPT2, and SPT2 were excluded from analyses B to E as they behaved as outliers likely due to technical reasons. Sample names are abbreviated as in Table 1.

### 2.2 Pool library preparation and sequencing

We performed whole-genome sequencing of pooled DNA (pool-seq) to assess the genomic variation among samples. This method provides population-level allele frequencies by sequencing to a high depth a single-barcoded library prepared from a mixture (pool) of DNA of individuals from a population (Schlötterer, Tobler, Kofler, & Nolte, 2014). DNA pools were prepared by mixing equal amounts of DNA of 30 to 96 individuals collected in close spatial and temporal proximity. DNA pools were quantified using a Qubit Fluorometer (Thermo Fischer Scientific Inc) aiming to have at least 1.5 µg of DNA in 25-50 µL and were submitted to the SNP&SEQ Technology Platform in Uppsala, Sweden, for library preparation and high-throughput sequencing. A PCR-free Illumina TruSeq library with an insert size of 350 base pairs (bp) was prepared for most pools, except for the ones extracted with Chelex, for which a Splinted Ligation Adapter Tagging (SPLAT) library was used instead, as it is aimed for single-stranded DNA (Raine, Manlig, Wahlberg, Syvänen, & Nordlund, 2017). Paired-end short reads (2×150 bp) were generated using an Illumina NovaSeq sequencer and S4 flow cells.

### 2.3 Read mapping and variant calling

Low quality bases, sequencing adapters, and short reads were removed from the raw reads using *Trimmomatic* v.0.36 (Bolger, Lohse, & Usadel, 2014). Clean reads were mapped against the *Trachurus trachurus* genome assembly (Accession: GCA_905171665.1, (Genner & Collins, 2022)) using *bwa-mem* 0.7.17 (Li, 2013). A pilot examination of read mapping statistics indicated that the temporal replicates from northern and southern Portugal (NPT2, SPT2) might be affected by technical artefacts (Figure S2). Hence, these samples were excluded from genome-wide analyses, but were included in analyses focused on specific loci (e.g., in the examination of allele frequency patterns).

Variant calling was performed using the algorithm *UnifiedGenotyper* of *GATK* v3.8 (McKenna et al., 2010). Biallelic SNPs were retained and various quality filters were applied to remove spurious markers (see Supporting Information for details, Figure S3). The resulting high-quality SNPs were used in further analyses. A summary of the data generation steps is shown in Figure S4.

### 2.4 Population genetic structure and genetic diversity

We assessed population structure using pairwise pool-*F*_ST_ and principal components analysis (PCA). We computed the pool-*F*_ST_ 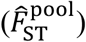 statistic using the R package *poolfstat* (Hivert, Leblois, Petit, Gautier, & Vitalis, 2018), which is equivalent to the Weir & Cockerham (1984) *F*_ST_ and accounts for random chromosome sampling in pool-seq.

To evaluate whether neutral or selective processes better explain the observed patterns of genetic differentiation, we separately performed PCA on two SNP subsets, one of undifferentiated (i.e., presumably neutral) and the other of highly differentiated markers (outliers, assumed to have been subject to selection). These marker sets were chosen based on the empirical distribution of allele frequencies and standard deviation (SD) cut-off values (Figure S5) (see Supporting Information for details). To reduce redundancy and physical linkage among SNPs, we retained, at most, one marker every 1 kb in the undifferentiated marker set, and one marker every 10 kb in the differentiated set, as it is expected that linkage is more pronounced in regions under selection. PCA was performed using the R package *prcomp*. In a pilot analysis one sample from the west of Ireland (WIE2) appeared as an outlier (Figure S6). In the absence of any plausible biological reason for this observation, this sample was removed.

To examine the genome-wide variation of genetic diversity in each pool, we calculated nucleotide diversity (π) per pool in 10 kb-sliding windows with a step size of 2 kb using *PoPoolation 1*.*2*.*2* (Kofler et al., 2011) (see Supporting Information for details). Plotting and statistical testing was performed using the R environment (R Core Development Team, 2021).

Additionally, we evaluated whether population structure followed an isolation-by-distance pattern with a Mantel test and 9999 permutations using the R package *ade4* (Dray & Dufour, 2007). We compared the linearized genetic distances, calculated with the formula linearized-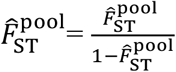 (François Rousset, 1997), and the geographic distances, estimated as the least-coast oceanic distance in kilometers (km) considering land as barrier using the R package *CartDist* (Stanley & Jeffery, 2017).

### 2.5 Detection of loci under selection

We applied effective coverage correction (*n*_eff_) to the raw read counts, in order to account for random variation of read coverage and chromosome sampling during pooling and sequencing (Bergland, Behrman, O’Brien, Schmidt, & Petrov, 2014; Feder, Petrov, & Bergland, 2012; Kolaczkowski, Kern, Holloway, & Begun, 2011) (see Supporting Information for details). Pool allele frequencies were then computed using a custom python script.

To identify genomic regions with elevated differentiation respect to the genomic background, we calculated the absolute delta allele frequency (dAF) per SNP between paired contrasts of grouped pools, as *dAF = absolute(meanAF(group1) – meanAF(group2)*. The contrasts evaluated were established based on geographic closeness, PCA clustering, and biological knowledge (Table S3). We also calculated the moving average of dAF values in 100-SNP windows to identify regions with consistent differentiation across nearby markers, while ruling out single SNPs that may be caused by random effects of pool-seq. We further explored the allele frequency patterns of the most differentiated SNPs at each locus and contrast for all the 12 pools. All the analyses were performed using R and plotting was done with the R package *ggplot2* (Wickham, 2016).

### 2.6 Validation of informative markers for genetic stock assessment

To identify a panel of highly informative SNPs for genetic stock identification and to validate pool-seq findings, we obtained the genotypes of 160 individuals (20 fish each from eight locations) in 100 SNPs (Table S4). The 100-SNPs panel consisted of 24 neutral markers and 76 putatively adaptive markers (see Supporting Information for details). The split of adaptive markers, in terms of observed association, was: North Sea (n=28), north-south break (n=13), west of Ireland (n=14), Alboran Sea (n=13), southern Portugal (n=4), north Africa (n=4). Three to four individuals per location were genotyped twice to assess genotyping error rate. DNA extraction and SNP genotyping were undertaken by IdentiGEN, Ireland, using their IdentiSNP genotyping assay chemistry. The protocol utilises target specific primers and universal hydrolysis probes. Following an end-point PCR reaction, different genotypes are detected using a fluorescence reader.

Based on individual allele frequencies, we undertook a preliminary analysis of population structure among the eight genotyped fish aggregations. It should be noted that sample sizes were small and therefore the results of the population analyses should be viewed as preliminary until further large-scale screening is undertaken. Only individuals and markers with >80% genotyping success were retained. Deviations from Hardy–Weinberg equilibrium (HWE) and linkage disequilibrium (LD) were assessed with *Genepop* 4.2 (Rousset, 2008).

Six SNPs had indication of deviation from HWE, two markers (12_3119866 and 17_972744) were not polymorphic and one had evident scoring errors (24_5252083), thus these nine markers were excluded. *Microsatellite Analyzer* (*MSA*) 4.05 was used to calculate pairwise *F*_ST_ estimates (Dieringer & Schlötterer, 2003). In all cases with multiple tests, significance levels were adjusted using the sequential Bonferroni technique (Rice, 1989). PCA was performed using the R function *prcomp*.

We estimated individual admixture coefficients for a given number of ancestral source populations (*K*) using the *sNMF* algorithm (Frichot, Mathieu, Trouillon, Bouchard, & François, 2014a) of the R package *LEA* (Frichot, Francois, & Caye, 2017). We tested *K* = 1 to 5, with 10 repetitions and 200 iterations. The most likely *K* corresponds to the value where the cross-entropy criterion (metric that evaluates the error of the ancestry prediction) plateaus or increases (Frichot, Mathieu, Trouillon, Bouchard, & François, 2014b). We plotted the average admixture proportions per population sample over a map using the R packages *ggplot* (Wickham, 2016) and *ggOceanMaps* (Vihtakari, 2020).

### 2.7 Characterization of a putative inversion on chromosome 21

To assess and compare the genetic diversity and spatial distribution of haplotypes of the putative inversion on chromosome (chr) 21, we extracted the individual genotypes of 12 diagnostic SNPs within the inversion from the 100-SNP dataset (Figure S7). We performed a PCA with the R function *prcomp* to identify the inversion genotype of each individual. Individuals were assigned to a haplotype group using the first two eigenvectors of the PCA and the k-means clustering algorithm implemented in the R function *kmeans*. We calculated observed heterozygosity for each of the PCA clusters, with the expectation that the middle cluster, presumably corresponding to inversion-level heterozygotes, will have the highest heterozygosity. These analyses and correspondent graphics were performed using R.

### 2.8 Genome-Environment Association

To identify which environmental variables are strongly correlated with adaptive genetic variation and local adaptation, we tested genome-environment association using redundancy analysis (RDA). We retrieved environmental data from *Bio-Oracle* v.2.1 (Assis et al., 2018; Tyberghein et al., 2012) for each sampled location using the *R* package *sdmpredictors* (Bosch, 2020). Data corresponded to mean bottom depth layers of minimum, mean, maximum, and range values of sea water temperature (°C), sea water salinity (PSS), and dissolved oxygen concentration (µmol/m^3^) for a 14-year period (from 2000 to 2014) and a spatial resolution of 0.25 arcdegrees. The data was derived from a numerical model developed by the Global Ocean Biogeochemistry non-assimilative Hindcast (PISCES) (Copernicus Programme of the European Union, n.d.).

Environmental data per location were standardized to zero mean and unit variance, and collinear variables were removed (see Supporting Information for details, Figure S8). To run RDA, we used the R package *vegan* (Dixon, 2003), the non-redundant environmental dataset, and the pool-allele frequencies of SNPs in outlier loci found with genome scans. The significance of the RDA model and environmental variables was assessed with an analysis of variance (ANOVA) using 1000 permutations.

### 2.9 Functional annotation of gene models

The gene models of the Atlantic horse mackerel genome were developed by Ensembl (Howe et al., 2021) and become available in an Ensembl Rapid release in March 2021 (Ensembl, 2021b). However, at the time this research was conducted, the gene models lacked gene symbols (names) and GO terms (only Ensembl gene IDs were available). Given that these annotations are relevant for interpretation of results, we ran the functional gene annotation pipeline developed by the National Bioinformatics Infrastructure Sweden (NBIS) (Binzer-Panchal, Dainat, & Soler, 2021) to retrieve this information (see Supporting Information for details).

Additionally, for the top 2% most differentiated SNPs within each divergent genomic region detected with genome scans, we annotated the closest overlapping gene (up to ± 40 kb) and the variant effect prediction (e.g., missense, synonymous, upstream, downstream, intergenic) using snpEff v.4.1 (Cingolani et al., 2012).

## 3. Results

### 3.1 Sequencing and variant calling

We generated pooled DNA whole-genome sequence data of 12 Atlantic horse mackerel aggregations sampled across the species’ range in the east Atlantic and the western Mediterranean Sea, Alboran Sea (Figure 1A, Table 1). Temporal replicates of four locations were collected one year apart, so the temporal stability of their genetic composition could be investigated (west of Ireland WIE1-WIE2, North Sea NOS1-NOS2, northern Portugal NPT1-NPT2, and southern Portugal SPT1-SPT2). Our sequencing effort yielded 490-764 million high-quality reads per pool. 98.2-99.2% of reads aligned to the genome assembly of *T. trachurus* (Genner & Collins, 2022), and each pool had a mean depth of coverage between 25.7x and 46.3x (Table S5). After variant calling with GATK, a total of ∼12.8 million biallelic SNPs passed quality filters and were used in the population analysis.

**Table 1.**
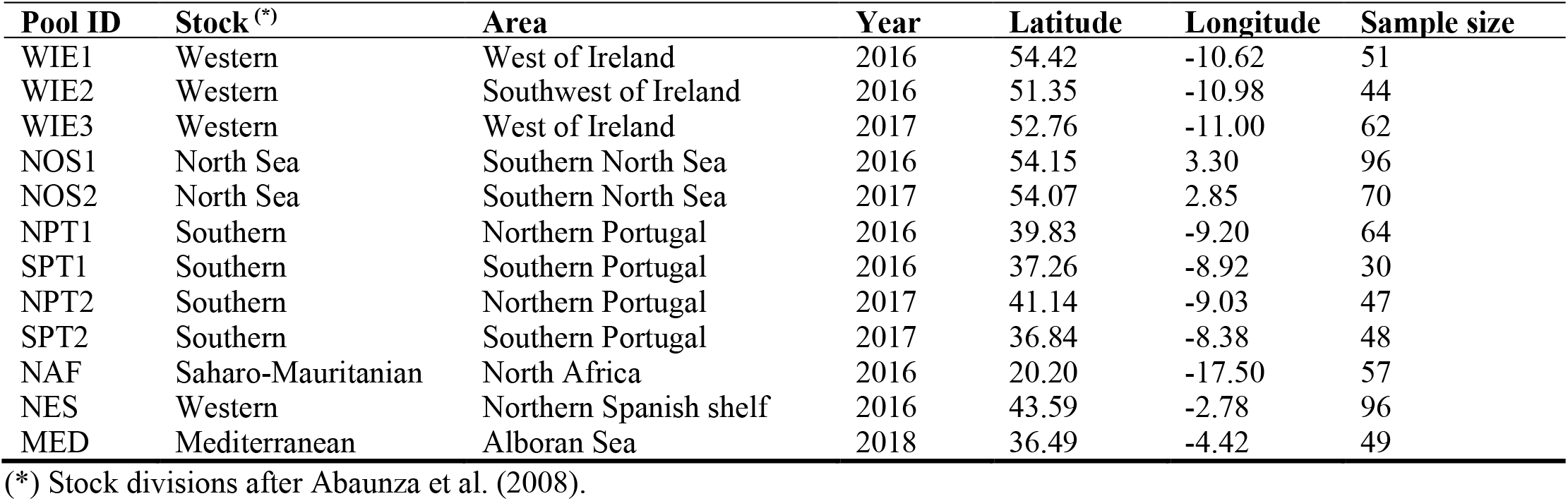
Collection details of the 12 Atlantic horse mackerel samples included in this study.

### 3.2 Population genetic structure

Pairwise pool-*F*_ST_ estimates indicated low genome-wide differentiation among horse mackerel populations (global mean pool-*F*_ST_=0.007, range 0.001-0.012) (Figure 1B), but revealed four subtle population structure patterns that were also supported by genome scans. First, the western Mediterranean Sea (MED) differentiates from Atlantic populations (mean pool-*F*_ST_=0.011). Second, Atlantic populations differ along a latitudinal cline that distinguishes “northern” or “southern” groups respect to a pronounced genetic break near Lisbon, Portugal (∼38.7-39.0°N). Third, populations within the “northern” group (North Sea– NOS1, NOS2, west of Ireland–WIE1, WIE3, northern Spanish Shelf–NES, northern Portugal–NPT1) are genetically more similar to each other (mean pool-*F*_ST_=0.004) than populations within the “southern” group (southern Portugal–SPT1 and north Africa–NAF) (mean pool-*F*_ST_=0.006). Forth, the two samples from the North Sea (NOS1 and NOS2) show the highest genetic similarity (pool-*F*_ST_=0.001) of all and are distinguished from other “northern” populations.

While genetic differences within the “northern” group are small, they are not negligible (pool-*F*_ST_ range 0.001-0.006) (Figure S9). A Mantel test assessing the degree of correlation between genetic and geographic distances (*H*o: no association) between pairs of populations indicated that subtle genetic differences within the northern group follow an isolation-by-distance (IBD) pattern, suggesting that they might be shaped by effective gene flow between neighbouring areas (Mantel test *p*-value=0.015; linear regression *R*.*adjusted*^2^ = 0.9, *p*-value = 1.1^-13^) (Figure 1C).

We further examined whether population structure patterns were driven primarily by neutral or selective processes by separately performing PCA for two SNP subsets, of undifferentiated or highly differentiated markers. These marker sets were chosen based on the standard deviation in allele frequencies across populations (Figure S5). The undifferentiated markers were considered “neutral” (61,543 SNPs) (Figure 1D, S8A), and the highly differentiated markers were assumed under selection (818 SNPs) (Figure 1E, S8B). In both PCA plots the separation between the “northern” and “southern” samples was evident, although the composition of the clusters slightly varied depending on the marker set. In both cases the “northern” group appeared as a tight cluster of genetically similar samples mainly from the North Sea, west of Ireland, the northern Spanish shelf, and northern Portugal. In stark contrast, the “southern” samples were dissimilar to each other. In the PCA with neutral markers, the first two axes explained 30.6% of the genetic variance. The mid-range samples from the Mediterranean Sea (MED) and southern Portugal (SPT1) were the most genetically distinct, while the North Africa (NAF) sample was genetically similar to the “northern” group. The North Sea samples tightly clustered with others in the “northern” group, being almost undistinguishable. In contrast, in the PCA with selective makers, the first two axes explained twice as much genetic variance (60.8%). The Mediterranean Sea sample was genetically closer to the “northern” group, while the other two “southern” samples, from southern Portugal and North Africa, were the most differentiated. Notably, the two North Sea samples formed a distinct cluster separated from other “northern” samples.

Taken together, these results indicate that the separation between the western-most part of the Mediterranean Sea and Atlantic populations is largely driven by neutral processes, while the latitudinal genetic cline and separation of North Sea samples could be the result of selective processes.

### 3.3 Putative loci under selection

We performed whole genome scans based on the absolute difference in allele frequencies (dAF) for paired contrasts to identify outlier loci respect to the genomic background (presumed under selection). The contrasts were chosen based on PCA clustering patterns (Figure 1D, 1E), and the outlier regions were identified using a Bonferroni Z-score threshold of significance. Our analysis revealed a number of genomic regions with elevated differentiation for three contrasts: (i) the Mediterranean Sea vs. others, (ii) “northern” vs. “southern” groups, and (iii) the North Sea vs. others (Figures 2-4). A summary of the top 2% most differentiated SNPs per region, their closest gene, gene function, and relative position to genes (e.g., missense, synonymous, upstream, downstream, intergenic) inferred with snpEff are compiled in Table S6.

**Figure 2.**
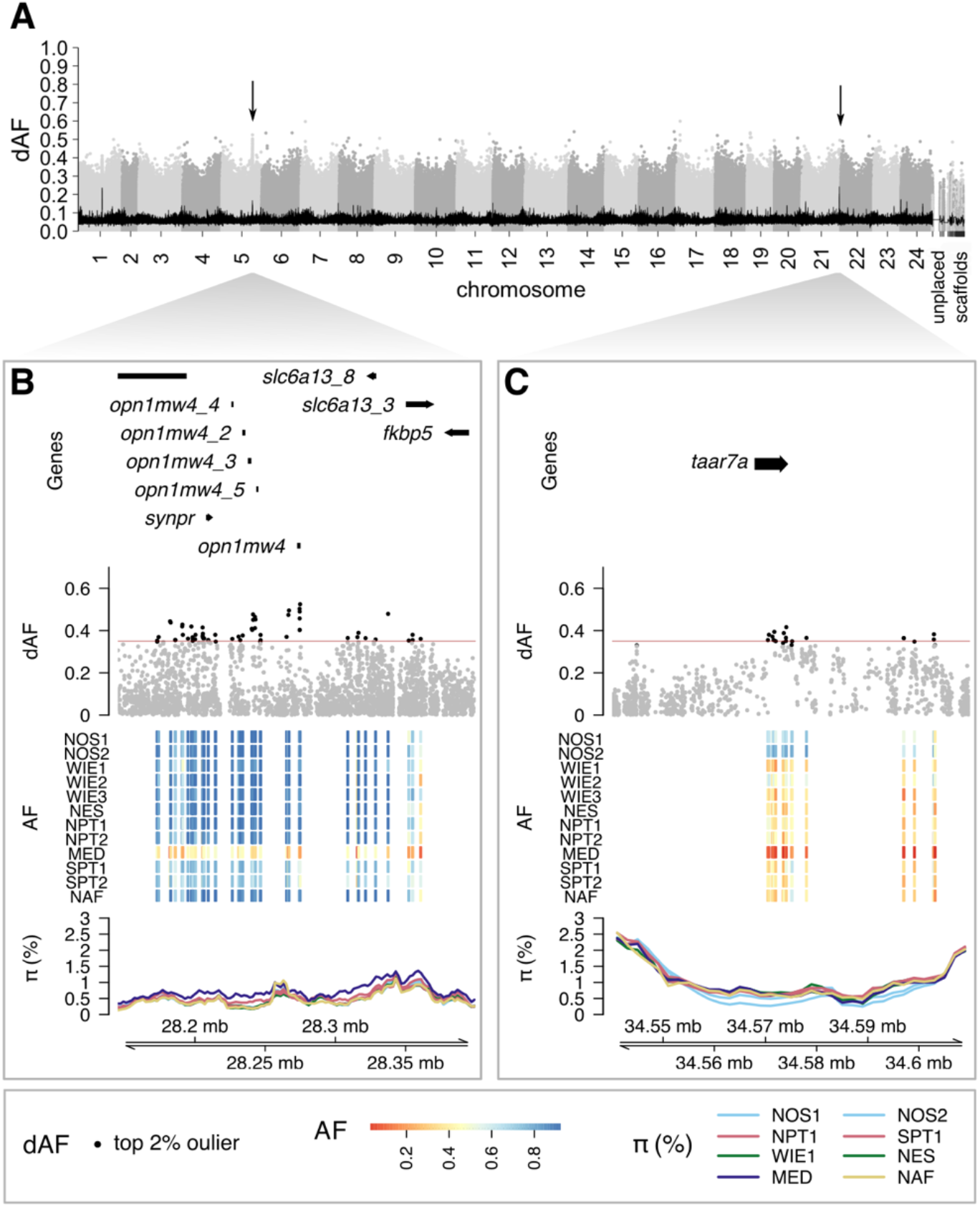
Two selective regions distinctive of the western Mediterranean Sea. (**A**) Manhattan plot depicting the dAF per SNP across the genome for the contrast between the Alboran Sea and all other samples. Each dot is a single SNP, and alternating gray tones were used to differentiate SNPs in consecutive chromosomes. The black line is the rolling average of dAF over 100 SNPs. Regions of interest are indicated with an arrow. (**B-C**) Zoom-in plots for the signals on (**B**) chr 5 and (**C**) chr 21. Each zoom-in plot consists of four tracks: the first, a representation of the gene models; the second, the dAF of SNPs, with the top 2% most differentiated SNPs denoted in black; the third, pool minor allele frequency of the top 2% SNPs in a heatmap plot, in which each row is one sample and each column is one SNP; and the fourth, the percentage of nucleotide diversity (π) calculated in 10 kb windows with 2 kb step size along the region with a separate line for each sample, colored based on the ICES stock divisions. Sample names are abbreviated as in Table 1.

**Figure 3.**
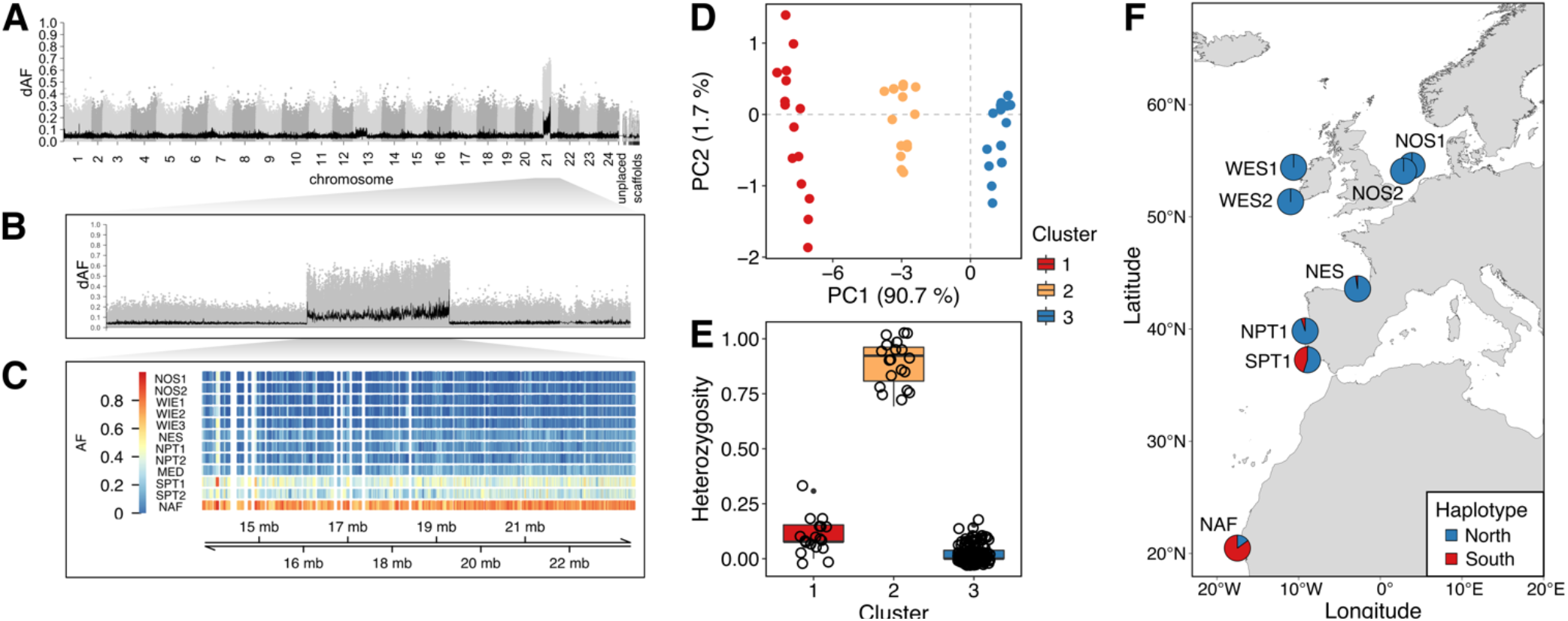
Putative chromosomal inversion on chromosome 21 underlies a latitudinal genetic cline. (**A**) Manhattan plot representing the absolute difference in pool-allele frequencies (dAF) of each SNP along the genome for the contrast between northern vs. southern groups (see Methods for details). The x-axis shows the genomic position of each SNP and the y-axis, its dAF value. SNPs in consecutive chromosomes are distinguished with alternating gray tones. The black line towards the bottom of the plot corresponds to the 100 SNPs-rolling average of dAF values. (**B**) Close-up Manhattan plot to chromosome 21. (**C**) Pool-allele frequency per sample (rows) of the top 2% markers (columns) within the putative inversion. (**D-F**) Analysis of inversion frequency based on the genotype of 15 SNPs screened in 160 individuals. (**D**) PCA plot showing individual clustering respect to the inversion genotype. (**E**) Individual observed heterozygosity in each PCA cluster. Clusters “1” and “3” correspond to homozygous “southern” and “northern” individuals, respectively, while cluster “2” are heterozygous individuals. (**F**) Map showing the geographic distribution of inversion haplotypes across sampled locations. Sample names are abbreviated as in Table 1.

**Figure 4.**
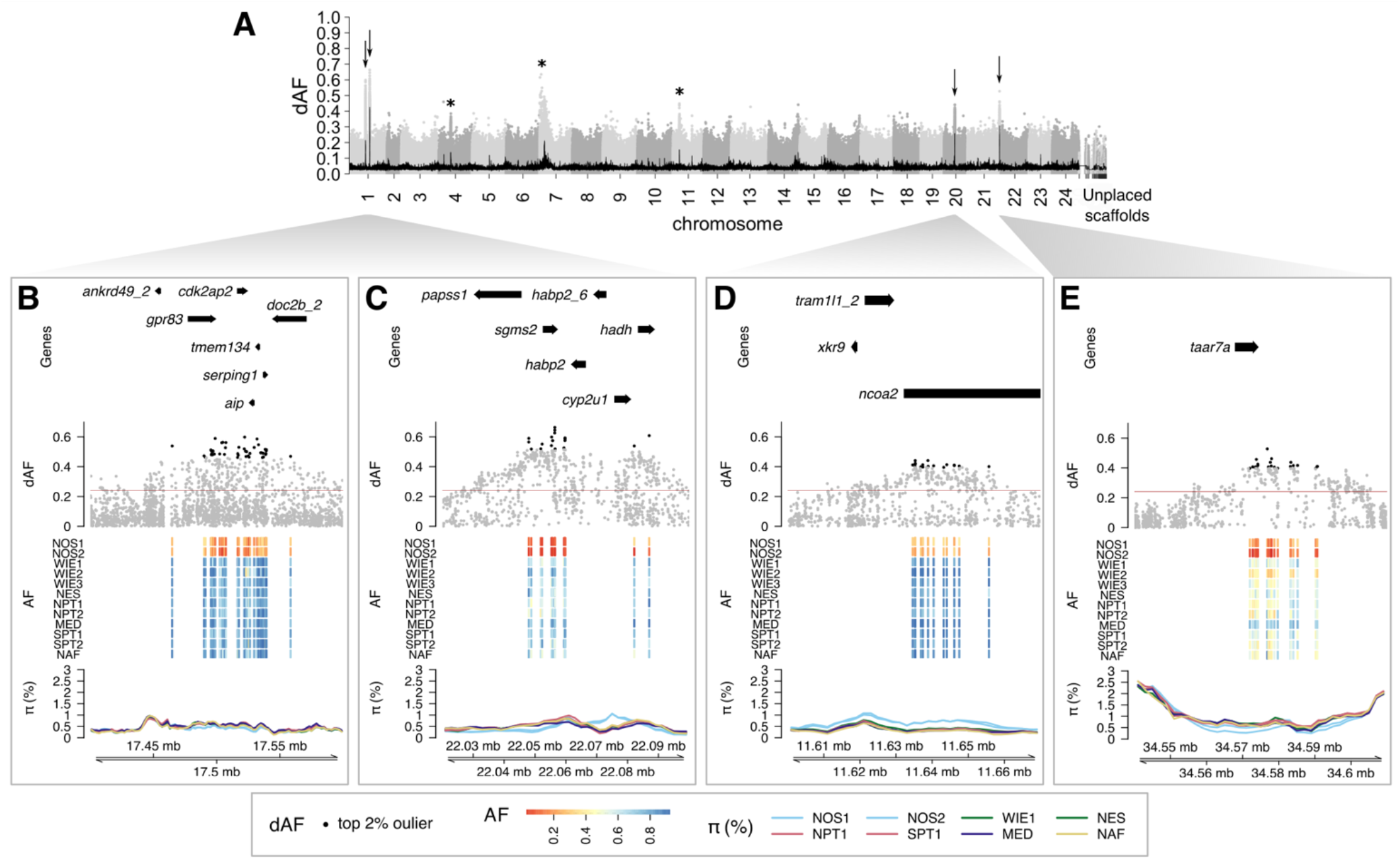
Genomic regions characteristic of the North Sea. (**A**) Manhattan plot representing the dAF of each SNP along the genome for the contrast between the North Sea and all other samples. Each dot is a single SNP, and alternating gray tones were used to differentiate SNPs in consecutive chromosomes. The black line towards the bottom is the rolling mean of dAF over 100 SNPs. (**B-E**) Close-up plots of the four most divergent regions (highlighted with arrows in A) on (**B-C**) chr 1, (**D**) chr 20, and (**E**) chr 21. Plots of the other differentiated regions on chr 4, 7 and 11 are shown in the Supplementary Figure S10 (highlighted with an asterisk in A). Each close-up plot consists of four tracks (from top to bottom): The first, illustrates gene models; the second, corresponds to the dAF of SNPs, in which the top 2% of markers are denoted in black. The horizontal red line indicates the Bonferroni Z-score threshold of significance; the third track, is a heatmap plot depicting the pool-allele frequency of the top 2% SNPs, where the rows are samples and the columns are SNPs; the fourth track, is the percentage of nucleotide diversity (π) for each sample calculated over 10 kb sliding windows with a step size of 2 kb. The color of each line indicates the designated ICES stock division of each pool prior to the HOMSIR project (Abaunza et al., 2008). Sample name abbreviations as in Table 1.

### Two outlier loci distinguish the western Mediterranean from Atlantic populations

The contrast between Mediterranean Sea and other samples revealed two distinctive regions at chromosomes (chr) 5 and 21 (Figure 2). In this case the signals of divergence were not as evident as in other contrasts given the overall higher genome-wide differentiation of this sample. Thus, we focused on the peaks in the rolling average of dAF values (Figure 2A), as it requires consistent allele frequency differences for the contrast of interest. We considered candidate genes those overlapping or located in the vicinity of the top 2% most differentiated SNPs in each locus. In the region on chr 5 (Figure 2B), the Mediterranean Sea sample had intermediate allele frequencies across loci and higher nucleotide diversity, while other samples tended to be fixed for one haplotype and showed lower nucleotide diversity. The top candidate gene in this region is *opn1mw4* (*green-sensitive opsin-4*). In the putative selection signal on chr 21 (Figure 2C), the Mediterranean sample had one haplotype close to fixation, the opposite haplotype was in high frequency in the North Sea, and intermediate frequencies were prevalent elsewhere. This region harbours a single gene, *taar7a* (*trace amine-associated receptor 7A*). The current knowledge of known functions of candidate genes are presented in Table S6.

### A presumed 9.9 Mb inversion distinguishes populations along a latitudinal cline

In the “northern” vs. “southern” contrast, we discovered that a single large region on chr 21 underpins the latitudinal cline with a genetic break along mid Portugal (Figure 3). This locus consists of a large block of SNPs with elevated dAF spanning 9.9 Mb (Figure 3A-B). The large size and abrupt drop in allele frequency differences towards the edges are characteristic of structural variants with suppressed recombination (e.g., inversions) that do not represent recent selective sweeps (Han et al. 2020). An analysis of the pool-allele frequencies of the top 2% most differentiated SNPs at this locus (Figure 3C) revealed striking allele frequency differences between the “northern” group (including the Mediterranean Sea) and North Africa, while intermediate allele frequencies were prevalent in southern Portugal.

To further characterize this structural variant, we leveraged genotype data of 160 individuals for 12 SNPs within the region. As expected for a chromosomal inversion, in a PCA plot individuals separated into three clusters depending on their genotype (Figure 3D), and individuals in the intermediate cluster exhibited the highest average observed heterozygosity (Figure 3E). Overall, the individual genotype data supported the observations made with pool-seq data regarding the spatial distribution of inversion haplotypes, although it offered greater resolution. For instance, it revealed that the “northern” haplotype dominates in the North Sea, west of Ireland, northern Spanish shelf, and northern Portugal, but also occurs at an intermediate frequency in southern Portugal. The “southern” haplotype is predominant in North Africa, and common in southern Portugal (Figure 1F). This putative inversion contains about 1,077 genes, making it difficult to infer which particular genes are under selection.

### A genetically distinct population in the North Sea

The contrast between the North Sea and other samples revealed seven genomic regions with elevated differentiation on chr 1, 4, 7, 11, 20, and 21 (Figure 4A). Further examination of allele frequencies of top SNPs per locus indicated that the North Sea temporal replicates were very similar genetically, and that their allele frequency patterns were distinctive of this location (Figure 4B-E, Figure S10). For instance, in the selection signals on chr 1 and 21, North Sea samples had variant alleles close to fixation and a slight reduction in nucleotide diversity (π) (Figure 4B, 4C, 4E). At the outlier loci on chr 4, 11 and 20, North Sea samples had predominantly intermediate allele frequencies, with similar (chr 4 and 11, Figure S10B, S10D) or higher (chr 20, Figure 4D) nucleotide diversity with respect to other pools. Additional divergent regions on chr 7 and 11 showed a similar but noisier pattern, presumably due to the presence of complex structural variants at these loci (Figure S10C, S10D).

The most striking signals were located on chr 1, followed by those on chr 20 and 21 (Figure 4). The signal on chr 1 encompasses two regions (chr1:17.4-17.6 Mb, chr1:22.0-22.1 Mb). In the first region (Figure 4B), the most differentiated SNPs are in the vicinity of the gene *gpr83* (G-protein coupled receptor 83). In the second region (Figure 4C), the top candidate gene is *sgms2* (*sphingomyelin synthase 2*). The signals on chr 20 (Figure 4D) and chr 21 (Figure 4E) contain a single gene each, *ncoa2* (*nuclear receptor coactivator 2*) and *taar7a* (*trace amine-associated receptor 7A*). Note that the signal on chr 21 is the same as the one noted in the Mediterranean sample (see above). Additional signals on chr 4, 7 and 11 (Figure S10) were not as clear as the ones just described (e.g., smaller allele frequency differences and/or inconsistent allele patterns). Therefore, it was difficult to identify candidate genes for these regions. The current knowledge concerning the function of candidate genes in the differentiated genomic regions is summarized in the Table S6.

### 3.4 Genome-Environment Association

To identify the environmental variables likely driving adaptive genetic variation, we retrieved seawater-related environmental variables (temperature, dissolved oxygen and salinity) and examined genome-environment association (GEA) based on redundancy analysis (RDA) (Figure 5A). A total of 210,568 SNPs within the most divergent genomic regions in the North Sea, Mediterranean Sea and northern vs. southern contrasts were used in this analysis.

**Figure 5.**
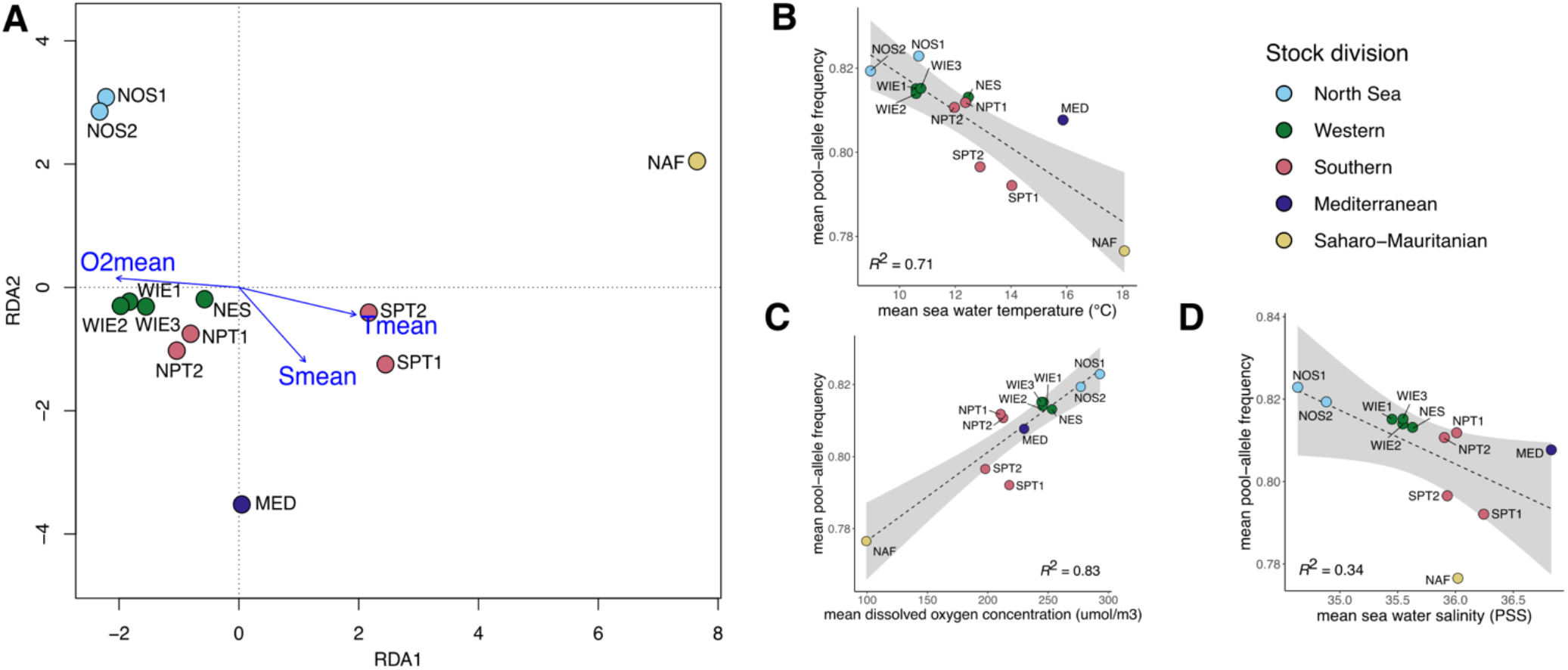
Association between putatively adaptive genetic variants and environmental variables. Analysis based on the pool-allele frequencies of 210,568 SNPs within the most divergent genomic regions in the North Sea and the Mediterranean Sea, and underlying the north-south genetic break. (**A**) Redundancy Analysis (RDA) plot showing the association between pool-allele frequencies of SNPs and three uncorrelated environmental variables: *T*_*mean*_, mean sea water temperature (°C); *O2*_*mean*_, mean dissolved oxygen concentration (µmol/m^3^); and *S*_*mean*_, mean sea water salinity (PSS). Blue arrows represent loadings of the environmental variables that contribute the most to the first two axes. Each dot represents a single pool sample, and its color indicates its ICES stock division. (**B-D**) Scatter plots showing the linear relationship (dotted line) and 95% confidence interval between average pool-allele frequencies and (**B**) seawater temperature, (**C**) mean dissolved oxygen concentration, and (**D**) mean seawater salinity (PSS). Sample name abbreviations, NOS1-2: southern North Sea, WIE1-3: western Ireland, NPT1-2: northern Portugal, SPT1-2: southern Portugal, MED: Mediterranean, NAF: North Africa.

Mean seawater dissolved oxygen concentration has the strongest association with genetic variation (*R*^*2*^ = 0.83), followed by mean seawater temperature (*R*^*2*^ = 0.71) and, to a lesser extent, with salinity (*R*^*2*^ = 0.34) (Figure 5B-D). Notably, the latitudinal genetic gradient discovered shows an inverse relationship with oxygen content, with the North Sea being the location with the highest oxygen levels (∼270-300 µmol/m^3^), almost three times the oxygen content in North Africa (∼100 µmol/m^3^) (Figure 5C). The two southernmost populations studied (in North Africa and southern Portugal)—which showed a high and intermediate frequency of the “southern” inversion haplotype, respectively—are from waters with higher mean temperatures (18°C), compared to locations north of Lisbon, in which the “northern” haplotype dominates. The North Sea has the coldest temperatures of all locations (< 10°C) (Figure 5B). In terms of salinity, as expected, the Mediterranean Sea has the highest salinity followed by southern Portugal, while the lowest salinity is observed in the North Sea (Figure 5D). That being said, the salinity range between the northern- and southernmost locations here analysed is minimal, between 35.0 and 36.5 PSS, which is considered a fully marine environment.

### 3.5 Validation of informative markers for genetic stock identification

To confirm the selection signals found with pool-seq data and develop a tool for genetic stock identification, we genotyped 160 individuals using 100 SNPs. Of these, 24 markers were considered neutral and 76 were among the most differentiated across contrasts (Figure S7). We found a strong correlation between allele frequencies calculated from individual genotypes and pool-seq data (mean *R*^2^ = 0.9 ± 0.1), supporting the findings of the pool-seq analysis (Figure S11). Of the 76 outlier loci, 72 had a genotyping success >80% (Table S7). After applying quality filters, the final dataset had 63 SNPs, while 157 out of 160 individuals had genotyping success >80%. Henceforth, this dataset will be referred to as the 63-SNP panel.

To minimise marker redundancy in the 63-SNP panel, we performed a linkage disequilibrium (LD) analysis for all loci and samples. As expected, significant LD was found between a number of SNPs located in close proximity on the same chromosome (Table S7). Though LD was not statistically significant in some cases (e.g., SNPs on chr 5), these were considered linked due to their physical closeness. To identify the most informative SNPs for discriminating the samples while reducing LD, we analyzed *F*_ST_ by marker and by population (Figure S12) for each genomic region. We retained the SNP with the highest average *F*_ST_ per linkage group (assumed to be the most informative), yielding a 17 SNP dataset comprising 155 out of 160 individuals with a genotyping success >80% (henceforth: the 17-SNP panel). Further analyses were conducted with both the 63-SNP and the 17-SNP datasets (individual genotypes are shown in Figure S13).

An examination of pairwise *F*_ST_ values indicated lack of significant genetic differences between the North Sea temporal replicates or between the two west of Ireland samples (Table S8). There was also no significant genetic differentiation between the individuals from the northern Spanish shelf, northern Portugal, and the two samples from the west of Ireland (Table S8).

To assess whether the same population structure is recovered with both the 63- and 17-SNP panels, we performed separate PCA and admixture analyses. We observed similar clustering patterns with both panels (Figure 6A, 6B), consisting of four main groups: (i) North Sea (NOS1, NOS2); (ii) west of Ireland (WIE1, WIE2), northern Spanish shelf (NES), and northern Portugal (NPT1); (iii) southern Portugal (SPT1); and (iv) north Africa (NAF). While the 63-SNP panel provided better discrimination between southern Portugal (SPT1) and North Africa (NAF), both marker sets separated most individuals from the North Sea, as well as the “northern” and “southern” groups. It should be noted that some individuals clustered in different groups from the ones expected, suggesting that gene flow occurs among geographic regions.

**Figure 6.**
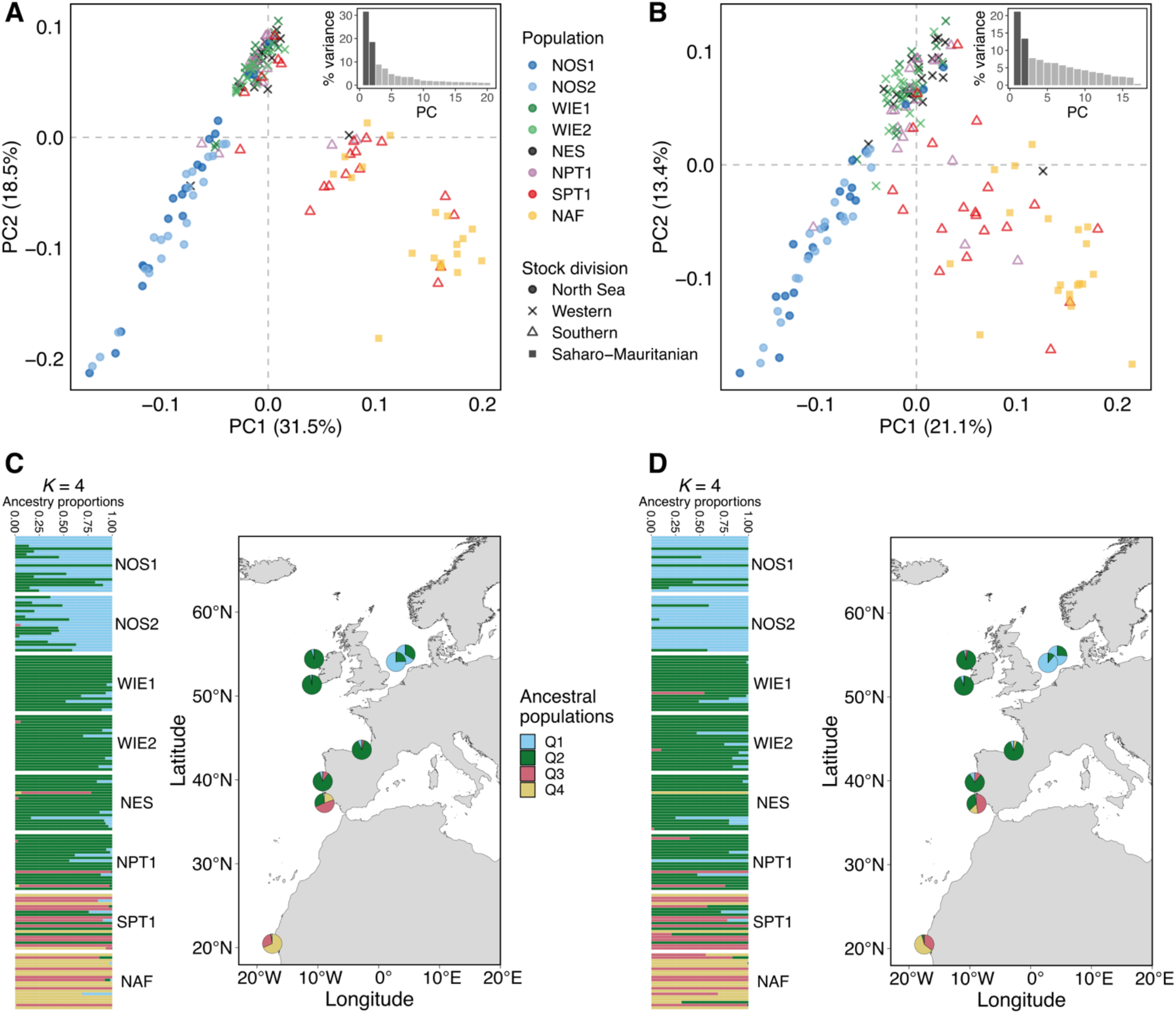
Population structure based on individual genotypes of reduced SNP panels. The two SNP datasets consisted on the genotype of 157 and 155 individuals screened using 63 and 17 markers, respectively. (**A-B**) Principal components analysis plot based on (**A**) the 63-SNP dataset, or (**B**) the 17-SNP dataset. Only the first two axes are shown. Each dot corresponds to an individual, the dot color indicates sampling location and the point shape, the designated stock based on the ICES stock divisions. Inset bar plots show the percentage (%) of genetic variance explained by each of the first nine principal components (PCs). (**C-D**) Admixture bar plot (left) and map showing the mean ancestry proportion per pool sample (right) for a number of ancestral populations (*K*), here equals to four, based on (**C**) the 63-SNP dataset, or (**D**) the 17-SNP dataset. In the admixture bar plot, each row is one individual, and the different colors per individual represent the probability of ancestry for a given *K*. Individuals collected in the same site are grouped as a block. Sample names are abbreviated as in Table 1.

An individual admixture analysis with both datasets supported the same four major genetic groups (Figure 6C, 6D) as the ones identified with PCA, showing various levels of mixing among them. For example, a few individuals in the North Sea (*n* = 3-4) had a high probability of originating from the western group (first-generation migrants). Similarly, southern Portugal had a number of individuals that appear to originate from the western group (*n* = 3) or from the African group (*n* = 3). In all four groups some individuals showed admixed ancestry, indicating that they are probably F1-hybrids or backcrosses between local and migrants. Overall, these results indicate that gene flow occurs more often between neighboring geographic areas, following an isolation-by-distance pattern (e.g., between the North Sea and the west of Ireland; between the northern Spanish shelf, northern Portugal and southern of Portugal; and between southern Portugal and north Africa).

## 4. Discussion

This study represents the most comprehensive assessment of the genomic variation of Atlantic horse mackerel populations to date (Figure 1). The combination of extensive geographic sampling and genomic screening derived from whole-genome sequencing, provided a powerful dataset that allowed us to discover genomic regions underlying population structure and local adaptation. Moreover, we developed a genetic tool that can distinguish the main biological groups.

### Population structure and temporal stability of genetic groups

Our genomic analysis indicated low genetic differentiation among horse mackerel populations inhabiting the area from the North Sea to north Africa (Figure 1A), more than 4,000 km of coastline (global mean pool-*F*_ST_=0.007, range: 0.001-0.012). This result is in agreement with previous studies using few genetic markers (Cimmaruta et al., 2008; Comesaña et al., 2008; Healey et al., 2020; Kasapidis & Magoulas, 2008), and implies that gene flow is pervasive. We infer that in the horse mackerel, since the larval stage lasts about one month (Rusell, 1976), gene flow could predominantly occur between neighbouring areas through passive transport of pelagic eggs and larvae by ocean currents. Moreover, the large population sizes of this species imply a minimal role of genetic drift in shaping patterns of genetic diversity.

Despite low differentiation at neutral loci, we uncovered patterns of population structure (Figure 1B, D, E) mainly at loci putatively under selection (Figures 2-4). Below, we will touch on the summary statistics results, and in the next section we will discuss the genome scan results.

Genetic differences between the Alboran Sea (the westernmost part of the Mediterranean) and the Atlantic occurred across the genome, implying that genetic drift contributed to the genetic differentiation. This separation was already proposed in earlier studies using body morphometrics, otolith shape, and parasitofauna (Abaunza et al., 2008). However, this is the first presentation of genetic data supporting this, as previous studies were inconclusive (Cimmaruta et al., 2008; Comesaña et al., 2008; Healey et al., 2020; Kasapidis & Magoulas, 2008). A Mediterranean-Atlantic divide has also been described in several marine species, but is not observed in all (Patarnello, Volckaert, & Castilho, 2007), likely because of differences in species’ life history or low resolution of the genetic markers used. Previous research based on parasite composition indicated that the Alboran Sea is a mixing area of horse mackerel from the Mediterranean and the Atlantic (Abaunza et al., 2008; Mattiucci et al., 2008). If this were the case, we would expect to see both inversion haplotypes at this location, but what we observed is that the northern haplotype prevails. This, in addition to the fact that none of the individuals collected were actively spawning (Table S1), let us infer that our sample is probably not representative of this area, and likely comes from a region with colder waters (e.g. Gulf of Lyon, Balearic Sea) (Figure S14). A possible neutral mechanism for reduced gene flow between Atlantic and western Mediterranean populations is the presence of retentive currents in the Almeria-Oran front, which have been proposed to act as barriers for gene flow for various marine species (Patarnello et al., 2007).

Our genomic data also revealed a hitherto undescribed latitudinal break off mid Portugal, near Lisbon (∼38.7-39.0°N), distinguishing populations from north and south of this area. The genetic separation was noticeable in pairwise-*F*_ST_ estimates, but more evident in the PCA based on outlier SNPs, suggesting a putative role of natural selection. Interestingly, the inferred location of the north-south genetic break coincides with a major biogeographical transition zone between temperate and subtropical waters off the coast of Portugal (Cunha, 2001; Santos et al., 2007). Moreover, a recent genetic study in the boarfish (*Capros aper*), a pelagic fish with similar distribution and life history as the horse mackerel, reported a comparable latitudinal pattern (Farrell, Carlsson, & Carlsson, 2016). Thus, it is possible that the environmental transition zone in Portugal is a major driver of population structuring of many marine species inhabiting this area, including the horse mackerel.

Within the “northern” group—North Sea, west of Ireland, the northern Spanish shelf (Bay of Biscay), and northern Portugal—genetic differences are small at both, neutral and selective markers (Figure 1D, E). Such genetic variability resembles an isolation-by-distance pattern (Mantel test *p*-value=0.015), which was also observed in a previous allozyme-based study (Cimmaruta et al., 2008). Thus, genetic differences between populations within the “northern” group seem to be largely determined by the effective gene flow via passive larval dispersal by ocean currents. The North Sea samples, though, were slightly more differentiated at outlier SNPs (Figure 1E), suggesting a putative role of natural selection.

In contrast, populations within the “southern” group—inhabiting waters in southern Portugal and North Africa, Mauritania—are genetically more differentiated between them and from all other populations. Interestingly, the population in the south of Portugal was different at both, neutral and selective markers, whereas the population from North Africa was more genetically distinct at outlier markers.

In summary, our genomic data indicates five possible genetic groups: (1) the western Mediterranean Sea; (2) the North Sea; (3) the west of Ireland, northern Spanish shelf (Bay of Biscay), and northern Portugal; (4) southern Portugal; and (5) north Africa, Mauritania. These groups seem to be temporally stable, as indicated by similar allele frequencies at outlier loci observed between temporal replicates (west of Ireland WIE1-WIE2, North Sea NOS1-NOS2, northern Portugal NPT1-NPT2, and southern Portugal SPT1-SPT2) (Figures 2-4).

### Genomic regions and environmental factors involved in local adaptation

Genome scans uncovered a number of regions with elevated differentiation that support the subtle patterns of population sub-structuring identified with summary statistics. The informative contrasts were: (i) the Mediterranean Sea vs. others, (ii) “northern” vs. “southern” groups, (iii) the North Sea vs. others. The divergent genomic regions varied in size (70-600 kb, and 9.9 Mb for a structural variant) and harboured numerous candidate genes.

The test for genome-environment association (GEA) indicated that genetic differences at putatively adaptive loci are primarily driven by differences in seawater oxygen content and temperature between locations (*R*^*2*^=0.83 and *R*^*2*^=0.71, respectively) (Figure 5). Notably, there was an inverse relationship between sea water oxygen, temperature, and the latitudinal cline, where the North Sea had the highest oxygen concentration (∼270-300 µmol/m^3^), almost three times the oxygen content in north Africa (∼100 µmol/m^3^). Moreover, the North Sea also exhibited the coldest seawater temperature of all locations analyzed (< 10°C), strikingly different from the two southernmost populations sampled (north Africa and south Portugal) (18°C).

We found that two outlier loci distinguish the western Mediterranean Sea, one on chr 5 and the other on chr 21 (Figure 2). The signal on chr 21 harbours a single gene, *taar7a* (*trace amine-associated receptor 7A*), which encodes a receptor involved in olfactory sensing of amines (Hashiguchi & Nishida, 2007). The signal on chr 5 includes three genes that encode proteins with crucial roles in synapsis (*cadps, synpr, slc6a13*), and five in-tandem copies of the green-sensitive opsin-4 (*opn1mw4*) gene. This gene organization (*opn1mw4* copies flanked by *synpr* and *slc6a13*) is highly conserved among several Clupeocephala species (e.g., Atlantic herring, zebrafish, medaka, stickleback (Lin, Wang, Li, & Wang, 2017; Nakamura et al., 2013)).

The top candidate gene at this locus is *opn1mw4*, a paralog of the *opn1mw* (RH2) gene, which encodes cone photopigments essential for vision of blue-green light. This gene contains two missense mutations (c.850G>A and c.670G>A, Figure S14A, B), one more divergent than the other (dAF=0.53 and dAF=0.40, respectively). The most divergent variant results in an amino acid change from alanine to tryptophan at residue 284 (p.Ala284Thr), and the less divergent variant, from valine to isoleucine at residue 224 (p.Val224Ile). While all four amino acids are hydrophobic, tryptophan is more hydrophobic and occupies a larger in volume than alanine, and the same is true for isoleucine in respect to valine, suggesting that these amino acid changes could potentially affect protein structure. Therefore, it is possible that the missense mutations in *opn1mw4* are causal, as affecting the structure of the photopigments could generate a shift in spectral sensitivity similar to the Phe261Tyr substitution in rhodopsin present in many fish species that live in brackish or freshwater (Hill et al., 2019).

A change in visual sensitivity could be an adaptive response to contrasting light environments between the Mediterranean and the Atlantic. In clear waters the blue-green light spectrum dominates, especially as depth increases, while in shallower or turbid waters red-shifted light conditions prevail. Comparative genomic studies indicate that fish living in marine habitats dominated by green-blue light (e.g., in the deep sea, the pelagic open ocean, or in coral reefs at night) commonly show gene copy expansions of *opn1mw* (RH2) respect to fish living in shallower (<30 m) or more turbid waters (Lin et al., 2017; Musilova & Cortesi, 2021). Indeed, the horse mackerel has five tandem copies of green-light opsins, which might relate to their preferred habitat (100–200 m depth) (Brunel et al., 2016). Seawater turbidity in the Mediterranean Sea is generally lower than in the Atlantic Ocean (Figure S14C), suggesting that more green light might be transmitted to deeper layers of the water column.

Taken together, the probable adaptive significance of the missense mutations in *opn1mw4* is that they might be involved in spectral fine-tuning of green-opsins to downwelling light detection under contrasting light conditions between the Mediterranean Sea and the Atlantic. The former is likely dominated by blue-green light conditions, while in the latter conditions are comparatively red-shifted. Visual adaptation could confer survival advantages related to feeding, recognition of conspecifics, and escape from predators.

We discovered that the latitudinal genetic cline is underlined by a large (9.9 Mb) putative chromosomal inversion on chr 21 (Figure 3). The northern haplotype shows high frequency in the “northern” group and in the western Mediterranean Sea, while the alternative haplotype dominates in north Africa, and occurs at an intermediate frequency south of Portugal. This chromosomal inversion harbours thousands of genes, the roles of which cannot be resolved without further studies. However, considering the strong genetic differentiation it drives along an important oceanographic transition zone, we infer this inversion may play a fundamental role in local adaptation to contrasting thermal regimes: a temperate (cooler) climate in the north, tropical (warmer) in the south, and variable/intermediate at the transition zone in mid Portugal. A possible explanation for the fact that the “northern” haplotype is prevalent in the western Mediterranean Sea is that this sample might actually spawn in a different location with cooler waters than where it was collected (near the Gibraltar strait, ∼16 °C). Within the Mediterranean Sea, comparable temperatures to those in the northern region (∼11-12 °C) occur in the winter season in coastal waters of the Balearic Sea near Catalonia (Spain), the Gulf of Lyon (France), and the Ligurian Sea (northeast Italy) (Pastor, Valiente, & Khodayar, 2020). Indeed, previous studies have reported spawning events of horse mackerel near Catalonia during the winter months (Andreu & Rodriguez-Roda, 1953; Planas & Vives, 1953). Given that the individuals collected in the Alboran Sea were not actively spawning, it is plausible that they come from a region further north.

We show that the population in the North Sea is genetically distinct. We reached this conclusion because seven genomic regions clearly distinguish samples from this location from all others, and the temporal replicates showed nearly identical allele frequencies at outlier loci. This observation adds to previous morphometric and parasite data suggesting that horse mackerel from the North Sea differs from nearby Atlantic populations (Abaunza et al., 2008).

Some of the top candidate genes likely involved in local adaptation to the North Sea are *gpr83, sgms2, ncoa2*, and *taar7a. gpr83* (G-protein coupled receptor 83) encodes a receptor that plays an important role in the regulation of energy metabolism, feeding, reward pathway, and stress/anxiety responses in mice, for which it has been linked with the control of body weight (Gomes et al., 2016; Lueptow, Devi, & Fakira, 2018). *ncoa2* (*nuclear receptor coactivator 2*) encodes a transcriptional coactivator for steroid receptors that is presumably involved in glucose metabolism regulation (Bateman et al., 2021). Previous experimental studies indicate that fish adapted to cold climates often have higher metabolic rates than those adapted to warm climates (Wang, Tan, Jiao, You, & Zhang, 2014; White, Alton, & Frappell, 2012). Therefore, we infer that selection may favor alleles that result in increased energy metabolism required for adaptation to the colder environment of the North Sea.

*sgms2* (*sphingomyelin synthase 2*) encodes a protein involved in the synthesis of sphingomyelin, a major component of cell and Golgi apparatus membrane. Previous studies indicate that this protein is crucial to maintain cell membrane structure and fluidity at low temperatures in fish (Wang et al., 2014; Windisch, Kathöver, Pörtner, Frickenhaus, & Lucassen, 2011). Thus, we propose that genetic variants in this gene could facilitate cold tolerance in the North Sea.

*taar7a* (*trace amine-associated receptor 7A*) codifies an olfactory receptor specific for sensing amines in vertebrates (Hashiguchi & Nishida, 2007; Hussain et al., 2009; Tessarolo et al., 2014; Yamamoto et al., 2010). Amines are behaviorally relevant odorants proposed to play a critical role in intra- and interspecific communication in, for example, sexual attraction or avoidance of predators or rotting food (Dewan, 2021). A previous study by Shi & Wang (2010) on two goatfish species with contrasting bottom habitat preferences (*Mullus surmuletus* and *Mullus barbatus*) reported significant differences in the morphology of chemoreceptors. Such differences are proposed to be associated with increased sensitivity to chemical stimuli in the species living in muddy and deeper waters, which has reduced visual capabilities. Among the locations in the east Atlantic included in this study, the North Sea has the highest water turbidity of all (Figure S14). Thus, natural selection may favor *taar7a* alleles that confer an enhanced sense of smell under the reduced visibility in the North Sea.

Other candidate genes distinctive of the North Sea might play critical roles in reproduction (spermatogenesis-*ankrd49*, gametogenesis-*spin1*, fertilization-*cdk2ap2*); development of bone (*papss1*), skeletal muscle (*spin1*) and skin (*ptk6*); cell processes (*aip*, apoptosis-*nol4*-*nol4l*, nutrient-dependant growth-*srms*); energy metabolism and body weight (glucose-*eno2*-*ncoa2*, lipids-*cyp2u1*, nitrogen-*slc38a3*, weight maintenance-*hadh*); nervous and olfactory system (neurotransmitter release, *doc2b*; synaptic transmission, *slc38a3*); immune response (*tmem134, serping1, clec4e*); and osmolarity regulation at low temperatures (*hsd11b1l*). The complete list of genes located within regions of genetic differentiation and their putative function are presented in Table S6.

Interestingly, the number of loci involved and their degree of genetic differentiation in the Atlantic horse mackerel are relatively small compared with those in the Atlantic herring (Han et al., 2020), but is rather intermediate to the Atlantic herring and the European eel, as the latter constitutes a single panmictic population (Enbody et al., 2021). We propose that the most important explanation for the differences in genetic structuring between these three species is related to their respective spawning strategies, since spawning and early development constitute the most sensitive period of life for a fish, characterized by high mortality (Dahlke, Wohlrab, Butzin, & Pörtner, 2020) and thus strong selection. The Atlantic herring is a demersal spawner that breeds close to the coast in areas with marked environmental differences between subpopulations as regards to temperature, salinity, depth and biotic conditions (plankton production, predators, etc.). It is also presumed to show prevalent homing behaviour, that is, individuals return for spawning to the locations where they were hatched. In contrast, the Atlantic horse mackerel is a benthopelagic fish that spawns at deeper waters, near the shelf edge (100-200 m deep), at not well-defined spawning areas. This implies that the environmental conditions in which breeding occurs are comparatively less diverse. On the other extreme, to the best of our knowledge, all European eels spawn in the Sargasso Sea under similar environmental conditions.

In summary, we revealed a number of highly divergent genomic regions encompassing genes that may be involved in adaptation to local environmental conditions in the North Sea, the western Mediterranean Sea, and along a climate-related latitudinal cline in the Atlantic. In addition, our results highlight the importance of chromosomal rearrangements for local adaptation with gene flow.

### Implications for fisheries management

The genetic-based groups identified here are largely in agreement with the current horse mackerel stock divisions, informed by the results of the 2000-2003 HOMSIR project (Abaunza et al., 2008). However, our data do not support the current definition of the “southern stock” in Portuguese waters (Figure S1), and of the southern boundary of the “western stock”.

Our genomic data indicates that the “southern stock” actually consists of two biological units (Figures 1, 3, 6), not one as previously assumed. Samples from northern Portugal (north of Lisbon, ∼38.7-39.0°N) appear to be genetically closer to the “western stock”, while samples from southern Portugal (south of Lisbon) form their own group but are genetically closer to the samples from the “Saharo-Mauritanian stock”, in North Africa. To confirm these findings and assess the spatial and temporal trends of mixing between these areas, further studies are required, including a finer geographic sampling and screening of informative genetic variants in a large number of individuals throughout this area.

We did not find significant genetic differences between northern Portugal (currently considered part of the “southern stock”), northern Spanish shelf (Bay of Biscay), and the west of Ireland, implying that the southern boundary of the “western stock” could possibly be extended down to the north of Portugal. However, the minute genetic differentiation does not exclude the possibility that isolation in an ecologically-relevant timescale of interest for fisheries management might occur (Hauser & Carvalho, 2008).

Finally, our genomic data support the consideration of the Mediterranean Sea as a separate stock, as proposed by the HOMSIR project (Abaunza et al., 2008). While a single sample from the westernmost part of the Mediterranean was studied, its genetic distinctiveness suffices to infer that the Mediterranean horse mackerel likely constitutes a separate population from those in the Atlantic. Wide-scale sampling within the Mediterranean Sea is required to further explore the population structure in this region.

This study identified a number of genetic markers (SNPs) that can be used as a genetic tool for fisheries stock assessment. A panel of only 63 markers suffice to identify the main genetic subdivisions. In fact, using a reduced panel of only 17 markers, it is possible to differentiate individuals collected in “the North Sea stock” and “western stock”, and from the “northern” (North Sea, west of Ireland, northern Spanish shelf, northern Portugal, and the Mediterranean) and the “southern” (southern Portugal, north Africa) groups with respect to the genetic break outside Lisbon. Thus, these markers can help, for instance, to elucidate the extent of mixing between the Western and North Sea stocks in the English Channel (ICES Divisions 7.e and 7.d) and in ICES area 4.a in the northern North Sea.

## Acknowledgements

Authors want to specially thank the members of the Northern Pelagic Working Group of the European Association of Fish Producers Organisation (EAPO) and the Pelagic Advisory Council (PelAC) for funding this study, and to Martin Pastoors, Maria Manuel Angélico, Finlay Burns, Gersom Costas, Cindy Van Damme, Cristina Nunes, Brendan O’Hea, Ciaran O’Donnell, Michael O’Malley, Jan Beintema and all IEO, IPMA, MI, MSS, WUR scientists and crew on the survey and commercial vessels involved in sampling. We also thank Jens Carlsson and the EU Atlas Project for providing the Mediterranean sample. The project was also financially supported by Vetenskapsrådet (2017-02907) and Knut and Alice Wallenberg Foundation (KAW 2016.0063). The National Genomics Infrastructure (NGI)/Uppsala Genome Center provided service in massive parallel sequencing and the computational infrastructure was provided by the Swedish National Infrastructure for Computing (SNIC) at UPPMAX partially funded by the Swedish Research Council through grant agreement no. 2018-05973. Thanks also to the National Bioinformatics Infrastructure Sweden (NBIS) for the development of the pipeline used for functional gene annotation. Thanks to Scott Campbell for enriching discussions.

## Data Accessibility

Sequence data are available in NCBI under the bioproject XX, Genbank accession XX Custom scripts are available in the Github repository: https://github.com/LeifAnderssonLab/2022_Horse_mackerel_popgen.

## Author contributions

EDF and LA conceived the study and obtained funds. APF-P performed the bioinformatic data analysis. EDF performed the genotype-based data analysis. MEP contributed to data analysis. CGS performed laboratory work. APF-P, EDF, and LA wrote the paper with contributions from all other authors. All authors approved the manuscript before submission.

## Competing interest statement

The authors declare no competing interest.

## Notes

### Competing Interest Statement

The authors have declared no competing interest.

### Summary of Updates

Typos, Supplementary Figures citation in text

