## Supplementary material for "The genomic basis and environmental correlates of local adaptation in the Atlantic horse mackerel (*Trachurus trachurus*)": Supp_info

#### Table of contents:


| | |
|---|---|
| **Extended Materials and Methods** | Page 2-7 |
| **Figure S1** | Page 8 |
| **Figure S2** | Page 8 |
| **Figure S3** | Page 9 |
| **Figure S4** | Page 10 |
| **Figure S5** | Page 10 |
| **Figure S6** | Page 11 |
| **Figure S7** | Page 12 |
| **Figure S8** | Page 13 |
| **Figure S9** | Page 13 |
| **Figure S10** | Page 14 |
| **Figure S11** | Page 15 |
| **Figure S12** | Page 16 |
| **Figure S13** | Page 17 |
| **Figure S14** | Page 18 |
| **Table S1** | Page 19 |
| **Table S2** | Page 20 |
| **Table S3** | Page 20 |
| **Table S4** | Page 21 |
| **Table S5** | Page 21 |
| **Table S6** | Page 22 (separate file) |
| **Table S7** | Page 22-23 |
| **Table S8** | Page 24 |
| **Supplementary References** | Page 25-28 |

### Extended Materials and Methods

#### Read mapping and variant calling

Raw sequence quality per pool was examined with *FastQC* v0.11.8 (Andrews, 2010), and a summary report for all pools was generated with *MultiQC* v.1.7 (Ewels, Magnusson, Lundin, & Käller, 2016). Low quality bases (Phred score < 15), Illumina adapters, and short reads (< 36 bp) were removed using *Trimmomatic* v.0.36 (Bolger, Lohse, & Usadel, 2014) (parameters: ILLUMINACLIP:adapters.fa:2:40:15:8:true SLIDINGWINDOW:4:15 LEADING:15 TRAILING:15 MINLEN:36).

Reads were mapped against the *T. trachurus* genome assembly (Accession: GCA_905171665.1, (Genner & Collins, 2022)) using *bwa-mem* 0.7.17 (Li, 2013) (default parameters). Read mapping quality statistics, including the number of aligned reads and the average read depth of coverage were generated with *QualiMap* v.2.2.1 (Okonechnikov, Conesa, & García-Alcalde, 2015). Prior to variant calling, mapped reads in BAM file format were sorted using *SAMtools* v.1.10 (Li et al., 2009), duplicate reads were marked and read groups were added, both with *Picard* v2.20.4 (Broad Institute, 2020), and an index file was generated for each BAM file using *SAMtools*.

A pilot examination of read mapping statistics suggested that the two temporal replicates from Portugal (NPT2, SPT2) might be affected by technical artefacts. They showed a significantly smaller mean coverage and shorter insert size (~245 bp vs. ~400-465 bp) than any other sample (Figure S2). Perhaps this relates to using a different DNA extraction method and library preparation, as their DNA was single-stranded. Hence, these samples were excluded from genome-wide analysis (e.g., principal components analysis and genome scans), but were included in analyses focused on specific loci (e.g., in the examination of allele frequency patterns).

Variant calling was performed using the algorithm *UnifiedGenotyper* implemented in *GATK* v3.8 (McKenna et al., 2010). The *GATK-UnifiedGenotyper* is a single base caller that simultaneously identifies Single Nucleotide Polymorphisms (SNPs) and small indels (insertions and deletions). Biallelic SNPs were extracted from the raw variant set and a series of filters were applied to keep the markers with the best quality. First, we performed hard-quality filtering by retaining SNPs that passed cut-off values set from the genome-wide distribution of GATK variant quality annotations. The filters applied were: FisherStrand (FS) > 60.0, StrandOddsRatio (SOR) > 3.0, RMSMappingQuality (MQ) < 40.0, MappingQualityRankSumTest (MQRankSum) < -12.5, and ReadPosRankSumTest (ReadPosRankSum) < -8.0 (for more details on the GATK quality annotations, see (Broad Institute, 2019)). Next, we retained SNPs with a genotype quality (GQ) greater than 20, allowed for a missing rate per locus of maximum 20%, kept loci with a minor allele count of at least 3 reads (MAC), and removed monomorphic loci with *BCFtools* v.1.10 (Li et al., 2009). Lastly, to exclude spurious SNPs in copy number variants and repetitive regions, which often show excessively high coverage, we applied a depth of coverage filter as follows. Using the *R* environment (R Core Development Team, 2021), we built a depth of coverage distribution per pool based on the read depth (DP) per SNP

(Figure S3). We separately evaluated three cut-off values (from the most to the least stringent filter): mean ± 1 standard deviation, mode ± ½ the mode, and between 20x and 300x (300x corresponds to three times the mean coverage across pools). In each case, we retained the SNPs that met the coverage thresholds for all pools, excluding three samples that appeared as outliers in the sequence quality assessment (NPT2, SPT2 and NAF). We chose the database resulting from the 20-300x filter because it retained a large number of loci while excluding those with extremely high depth. The resulting high-quality SNPs were used in further analysis. A schematic summary of the data generation steps is shown in Figure S4.

**Population genetic structure and genetic diversity**

We assessed the population structure of the Atlantic horse mackerel using pairwise $F_{ST}$ and principal components analysis (PCA). We computed the pool-$F_{ST}$ ($\hat{F}_{ST}^{pool}$) statistic for all population pairs based on the raw read counts per SNP and using the R package *poolfstat* (Hivert, Leblois, Petit, Gautier, & Vitalis, 2018). This statistic is equivalent to the Weir & Cockerham (1984) $F_{ST}$ and accounts for random chromosome sampling in pool-seq. The pool-$F_{ST}$ statistic ranges between 0 and 1, where a value of 0 indicates no genetic differentiation between populations, and a value of 1 means complete genetic differentiation.

To evaluate whether patterns of population structure were better explained by undifferentiated (assumed neutral) or highly differentiated markers (outliers, assumed selective), we generated two SNP datasets based on the empirical distribution of allele frequencies and standard deviation (SD) cut-off values (Figure S5). The undifferentiated marker set consisted of SNPs with allele frequencies close to the mean distribution (0.03 < allele frequency SD ≤ 0.09), while the differentiated set comprised outlier SNPs with allele frequency ≥ 0.2 SD from the mean. To minimize the presence of physically linked loci in each group of markers, we retained one SNP every 1 kb in the undifferentiated marker set, and one SNP every 10 kb in the differentiated set, as it is expected that linkage is more pronounced in regions of selection. PCA was performed separately for each dataset with the R package *prcomp.* In a pilot PCA one of the samples from the west of Ireland (WIE2) behaved as an extreme outlier (Figure S6). Since no biological reason can explain this behavior (likely this pool was a mixture of populations with contrasting genomic background), this sample was excluded from the analysis.

We examined the genetic diversity of each pool with estimates of nucleotide diversity ($\pi$) obtained with *PoPoolation 1.2.2* (Kofler et al., 2011). For this we generated a pileup file from each BAM file using *samtools v.1.10* (Li, 2011). As we focused our study on SNPs, we excluded indels and likely spurious SNPs around indels (± 5 bp). The read coverage of each pileup file was subsampled (without replacement) to a common value to account for potential biases due to random coverage variation among pools during sequencing (Kofler et al., 2011). This value was set to the minimum coverage required for a SNP to be retained, which in our case corresponded to the 5% quantile of the per pool coverage distribution (~20x, Figure S3). We required that SNPs had a coverage between 5-99% of the per pool coverage distribution, a minimum base and mapping quality of 20, and a minor allele count of 2 to be included in the analysis. Nucleotide diversity was calculated in 10 kbsliding windows with a step size of 2 kb, requiring that windows had a minimum coverage fraction of 0.5. Plotting and statistical testing was performed using the *R* environment (R Core Development Team, 2021).

To evaluate whether population structure followed an isolation-by-distance pattern, we performed a Mantel test with 9999 permutations implemented in the R package *ade4* (Dray & Dufour, 2007). For this, we compared the linearized genetic distances (pairwise pool-$F_{ST}$ values) calculated with the formula linearized-$\hat{F}_{ST}^{pool}$= $\frac{\hat{F}_{ST}^{pool}}{1-\hat{F}_{ST}^{pool}}$ (François Rousset, 1997), and the geographic distances estimated as the least-coast oceanic distance in kilometers (km) considering land as barrier using the *R* package *CartDist* (Stanley & Jeffery, 2017).

**Detection of loci under selection**
Before calculating allele frequencies per pool, we rescaled the raw read counts to the 'effective coverage' ($n_{eff}$) per SNP, which is an estimate of the number of chromosomes sampled adjusted by the read depth. This correction accounts for random variation of read coverage and chromosome sampling across pools during sequencing (Bergland, Behrman, O'Brien, Schmidt, & Petrov, 2014; Feder, Petrov, & Bergland, 2012; Kolaczkowski, Kern, Holloway, & Begun, 2011). We applied this correction to the raw read counts using a python script implementing the equation $n_{eff} = \frac{(n*RD)-1}{n+RD}$, where *RD* is the read depth and *n* is the number of chromosomes in a pool, which is equal to 2*N* (*N* = number of individuals in a pool) in a diploid organism. Population allele frequencies were then computed based on the $n_{eff}$ corrected read counts with a python script.

To identify regions of the genome with elevated differentiation with respect to the genomic background, interpreted as candidate regions under selection, we calculated the absolute delta allele frequency (dAF) per SNP between paired contrasts of single or grouped pools, as
*dAF = absolute(meanAF(group1) – meanAF(group2)*. The contrasts evaluated were established considering geographic closeness, PCA clustering patterns, and biological knowledge (see details in Table S3). We also calculated the moving (or rolling) average of dAF values in windows of 100 SNPs to identify regions with consistent differentiation across nearby markers, while ruling out single SNPs that could be influenced by random effects of pool-seq experiments. We further explored the allele frequency patterns of the most highly differentiated SNPs at each locus and contrast for all the 12 pools. All the analyses were performed using *R* and plotting was done with the *R* package *ggplot2* (Wickham, 2016).

**Validation of informative markers for genetic stock assessment**
To identify a reduced panel of highly informative SNPs for genetic stock identification, and to validate the pool-seq findings, we obtained the genotypes of 160 individuals (20 fish each from eight locations) in 100 of the most differentiated SNPs (Table S4).

The 100-SNPs panel was chosen as follows. First, we selected the most differentiated SNPs (dAF ≥ 0.35) from each divergent genomic region per contrast.

We set a higher dAF cut-off (in 0.5 increments) when more than 100 SNPs passed this threshold, until 10 SNPs remained. We required SNPs had a coverage ≥ 20x, base quality ≥ 20, mapping quality ≥ 20, were at least 10 bp from an indel, were more than 100 bp from repetitive sequences, and more than 1 kb from the closest informative SNP. We also retained SNPs that had alleles equally supported by forward and reverse reads (did not show strand bias), and that had enough flanking sequence for primer design (± 120 bp). We further assessed whether the quality of the flanking region was optimal (had good read support and no evidence of poor alignment), by visually inspecting the BAM files using the genome browser *IGV* (Robinson et al., 2011; Thorvaldsdóttir, Robinson, & Mesirov, 2013).

We additionally chose a set of undifferentiated SNPs. These markers were randomly selected from the chromosomes that were not informative in the main contrasts, and had to fulfill the same requirements as the outlier SNPs. The final split of loci per region in the 100-SNP panel was: North Sea (n = 28), north-south break (n = 13), west of Ireland (n = 14), Alboran Sea (n = 13), southern Portugal (n = 4), north Africa (n = 4), undifferentiated loci (n = 24). Three to four individuals per location were genotyped twice to assess genotyping error rate. DNA extraction and SNP genotyping were undertaken by IdentiGEN, Ireland, using their IdentiSNP genotyping assay chemistry. The protocol utilises target specific primers and universal hydrolysis probes. Following an end-point PCR reaction, different genotypes are detected using a fluorescence reader.

Based on the individual allele frequencies, we undertook a preliminary analysis of population structure among the eight individually-genotyped fish aggregations. It should be noted that sample sizes were small and therefore the results of the population analyses should be viewed as preliminary until further large-scale screening is undertaken. Only individuals and markers with >80% genotyping success were retained in the analyses. Deviations from Hardy–Weinberg equilibrium and linkage disequilibrium were assessed with *Genepop* 4.2 (F. Rousset, 2008)2008) (default settings). Six SNPs had indication of deviation from Hardy-Weinberg Equilibrium (HWE), two markers (12_3119866 and 17_972744) were not polymorphic and one had evident scoring errors (24_5252083), thus these nine markers were excluded. *Microsatellite Analyzer* (*MSA*) 4.05 was used to calculate pairwise $F_{ST}$ estimates (Dieringer & Schlötterer, 2003) (default settings). In all cases with multiple tests, significance levels were adjusted using the sequential Bonferroni technique (Rice, 1989)1989). PCA was performed using the *R* function *prcomp*.

We estimated the individual admixture coefficients respect to a given number of distinct populations (K) using the sNMF algorithm (Frichot, Mathieu, Trouillon, Bouchard, & François, 2014a) of the R package LEA (Frichot, Francois, & Caye, 2017). We tested K = 1 to 5, with 10 repetitions and 200 iterations. The most likely *K* corresponds to the value where the cross-entropy criterion (metric that evaluates the error of the ancestry prediction) plateaus or increases (Frichot, Mathieu, Trouillon, Bouchard, & François, 2014b). We plotted the average admixture proportions per population sample over a map using the *R* package *ggplot* (Wickham, 2016) and *ggOceanMaps* (Vihtakari, 2020).

**Characterization of a potential inversion in chromosome 21**
To assess the genetic diversity and spatial distribution of haplotypes of the putative inversion in chromosome (chr) 21, we extracted the individual genotypes of 12 diagnostic SNPs within the inversion from the 100-SNP dataset (Figure S7). To identify the inversion genotype of each individual, we performed a PCA with the *R* function *prcomp*. Individuals were assigned to each haplotype group using the first two eigenvectors of the PCA and the k-means clustering algorithm implemented in the *R* function *kmeans*. We calculated observed heterozygosity for the individuals in each of the PCA clusters, with the expectation that the middle cluster will have the highest heterozygosity of all. These analyses and correspondent graphics were performed using the *R* environment.

**Genome-Environment Association**
To identify which environmental variables may be associated with local adaptation, we performed a redundancy analysis (RDA) in the *R* environment. Using the *R* package *sdmpredictors* (Bosch, 2020), we collected environmental data for the 12 sampled horse mackerel location from a numerical model developed by the Global Ocean Biogeochemistry non assimilative Hindcast (PISCES) (Copernicus Programme of the European Union, n.d.), available through *Bio-Oracle* v.2.1 (Assis et al., 2018; Tyberghein et al., 2012). Environmental data consisted of mean bottom depth layers of minimum, mean, maximum, and range values of sea water temperature (°C), sea water salinity (PSS), and dissolved oxygen concentration ($\mu mol/m^3$), for a 14-year period (from 2000 to 2014) and a spatial resolution of 0.25 arcdegree.

Prior to RDA, the extracted environmental data per location were standardized to zero mean and unit variance, and the redundancy between variables was assessed (multicollinearity). We calculated the pairwise correlation coefficients with the function *pairs.panels* of the *R* package *psych* (Revelle, 2018) (Figure S8). When collinearity was high $| R^2 > 0.7 |$, we retained one of the variables based on biological/ecological knowledge (Forester, Lasky, Wagner, & Urban, 2018). We also checked collinearity while running RDA using the variance inflation factor (VIF) of the RDA model, where the variable with the highest VIF was removed until all variables had a VIF < 5 (Zuur, Ieno, & Elphick, 2010). To run RDA, we used the R package *vegan* (Dixon, 2003), the filtered environmental dataset, and the pool-allele frequencies of SNPs within differentiated genomic regions in all contrasts. The significance of the RDA model and environmental variables was assessed with an analysis of variance (ANOVA) using 1000 permutations.

**Functional annotation of gene models**
The gene models of the Atlantic horse mackerel genome were developed by Ensembl (Howe et al., 2021) and are available in the Ensembl Rapid release website since March 2021 (Ensembl, 2021b). However, at the time this research was conducted, the gene models lacked gene symbols (names) and GO terms (only had Ensembl gene IDs). This information is relevant to infer the potential functional effect of genetic variants of interest. To gather this data, we ran the functional gene annotation pipeline developed by the National Bioinformatics Infrastructure Sweden (NBIS) (Binzer-Panchal, Dainat, & Soler, 2021).

In brief, the pipeline starts by extracting the nucleotide sequence of each annotated gene from the genome sequence in FASTA format using the gene coordinates from the GFF annotation file. The nucleotide sequence per gene is then translated into amino acid sequence using *Another Gtf/Gff Analysis Toolkit* (AGAT) (Dainat, 2021). To infer which protein corresponds to each gene, the amino acid sequence is compared against a reference protein database using BLASTp (Altschul, Gish, Miller, Myers, & Lipman, 1990) (blast e-value = 1e-6). We used as reference the reviewed Vertebrates protein database available from UniProtKB/SwissProt on July 2021 (version 2021_03) (Bateman et al., 2021). Finally, the functional annotations (gene names and GO terms) of the matching protein are retrieved from InterPro databases (Blum et al., 2021) using the program InterProScan v.5.52-86.0 (Jones et al., 2014), and are assigned to their corresponding gene.

Additionally, for the top 2% most differentiated SNPs within each divergent genomic region detected with genome scans, we annotated the closest overlapping gene (up to $\pm$ 40 kb) and the variant effect prediction (e.g., missense, synonymous, upstream, downstream, intergenic) using snpEff v.4.1 (Cingolani et al., 2012).

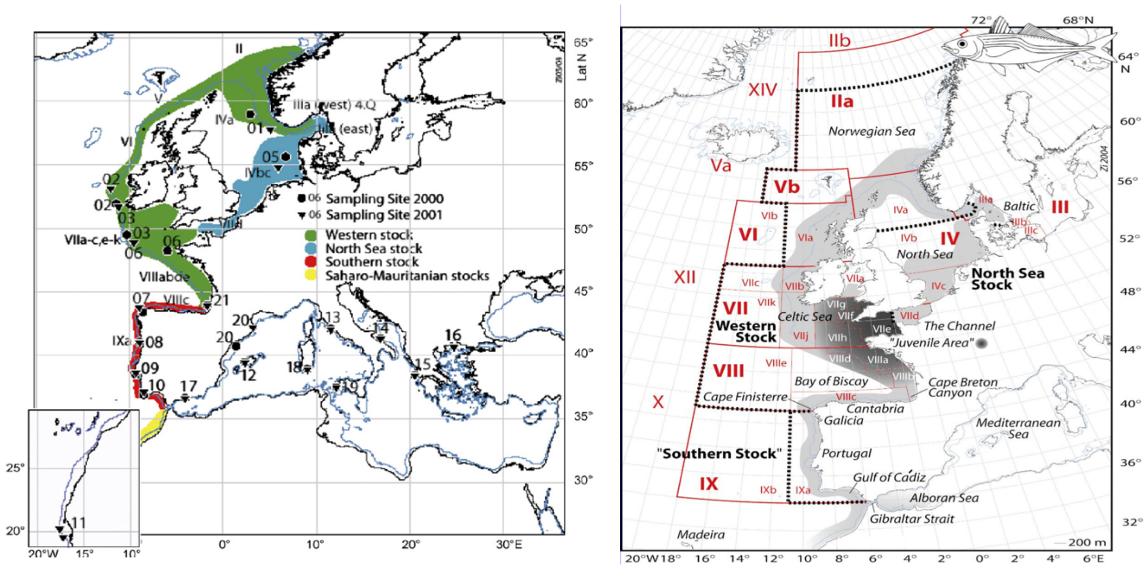


**Figure S1.** Northeast Atlantic horse mackerel stocks. (Left panel) Divisions prior to the HOMSIR project, image from Abaunza *et al*. (2008). (Right panel) Current stock divisions, after the HOMSIR project, image source (ICES, 2005).

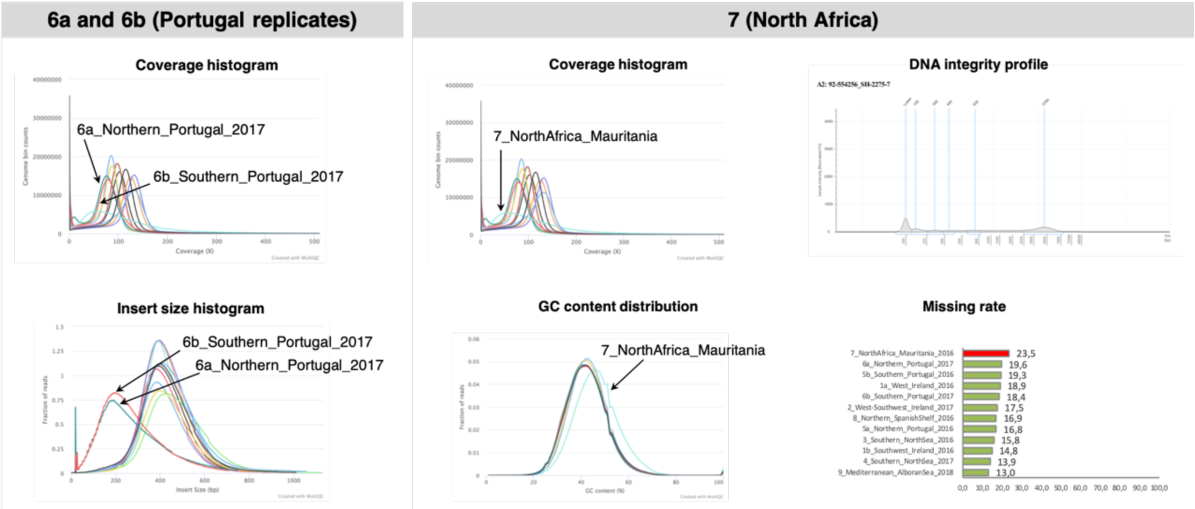


**Figure S2.** Read mapping statistics supporting that samples 6a, 6b, 7 were likely affected by technical artefacts. Plots obtained with *MultiQC*. (Left) Coverage and insert size distribution plots for the 12 samples, denoting the lines corresponding to samples 6a and 6b. (Right) Left, coverage and GC content distribution for all 12 samples, sample 7 is highlighted. Right, DNA integrity profile for the African sample and comparison of missing rate percentage for all 12 samples, the African sample is denoted in red.

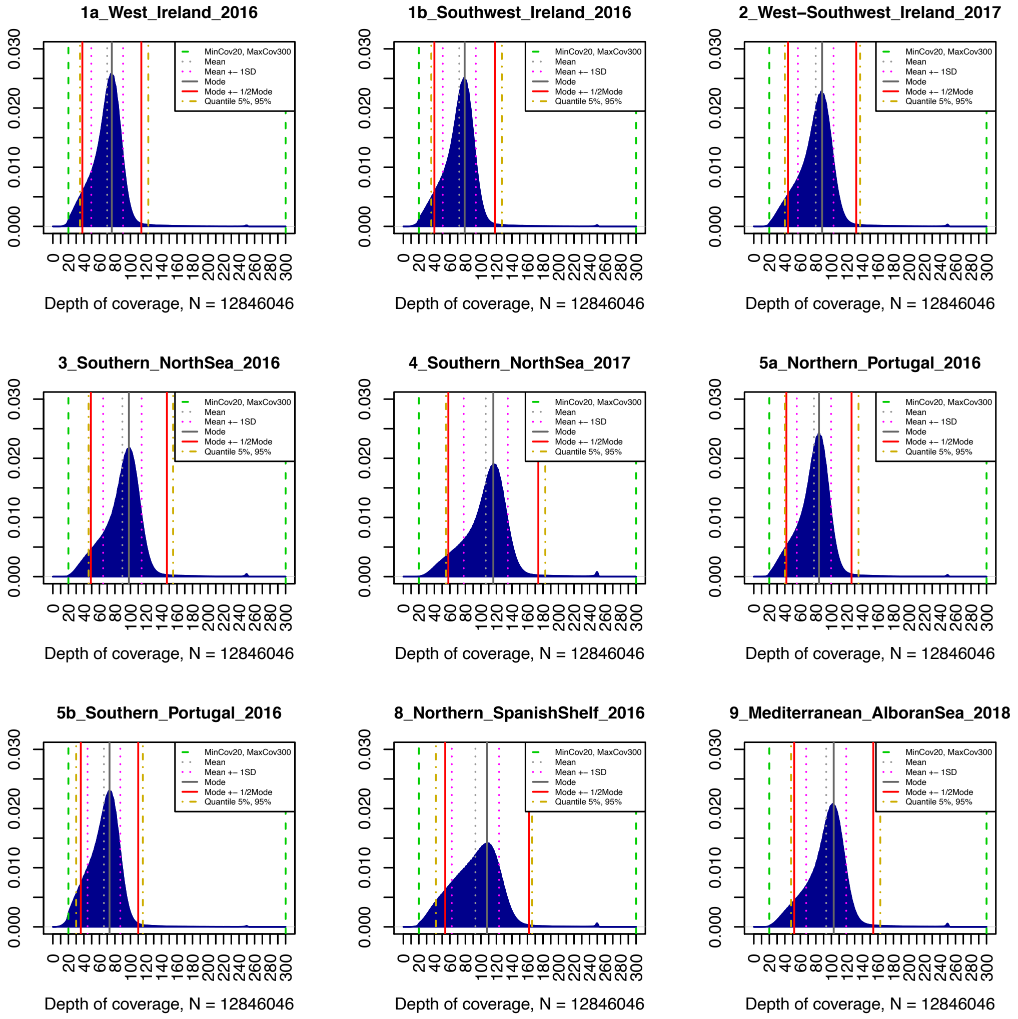


**Figure S3.** Depth of coverage distribution of horse mackerel pools based on the SNPs that passed quality filters (~12 million). The different vertical lines correspond to the various lower and upper depth of coverage cut-off values examined.

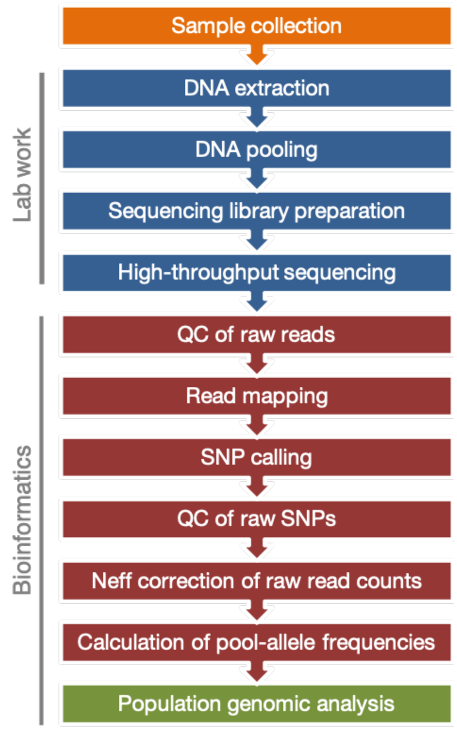


**Figure S4.** Schematic summary of steps followed for data generation.

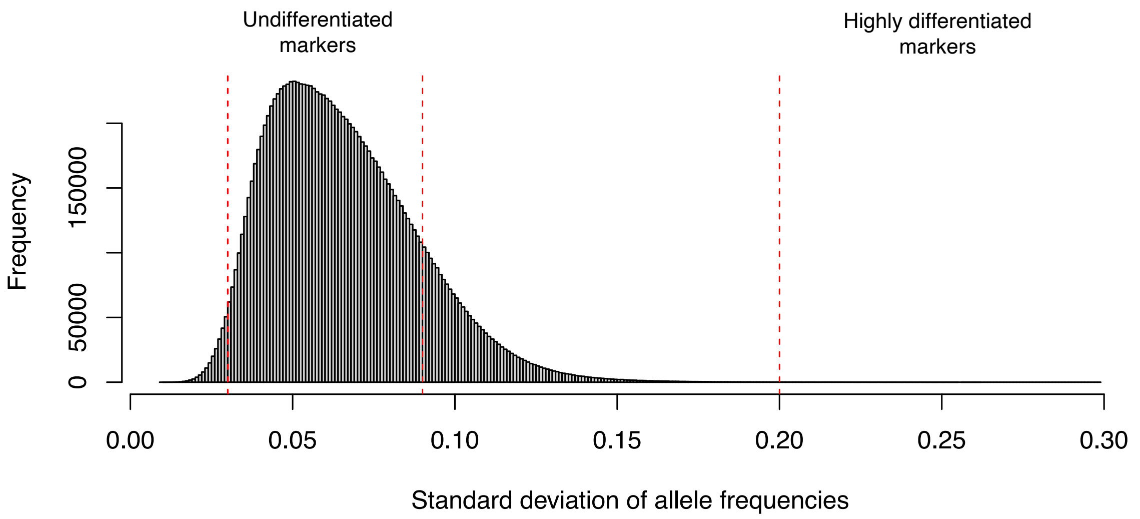


**Figure S5.** Histogram representing the allele frequency distribution for ~12 million SNPs. Vertical red dashed lines indicate the cut-off values to filter undifferentiated and highly differentiated markers.

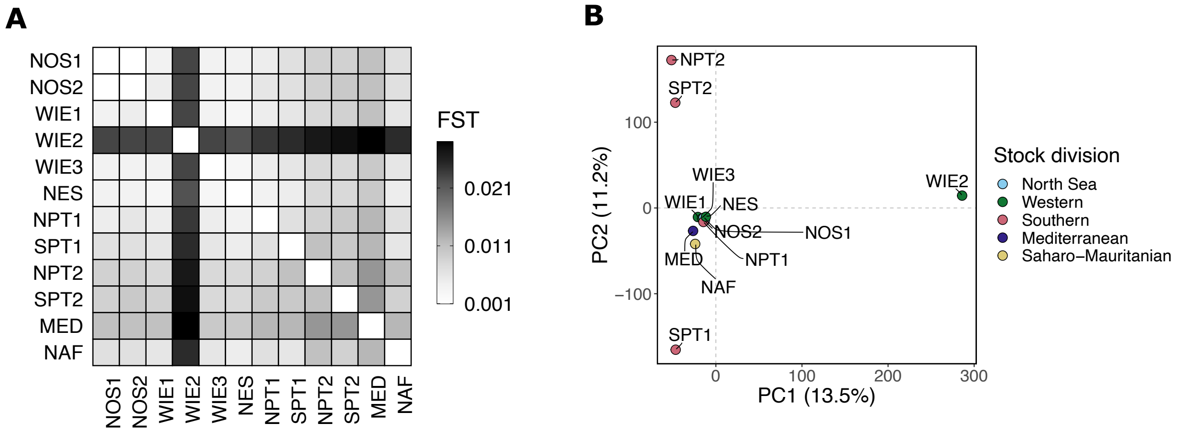


**Figure S6.** Pilot population structure analysis of the 12 Atlantic horse mackerel pools. (**A**) Pairwise $F_{ST}$. (**B**) PCA. The plots indicate that the sample WIE2 is an outlier.

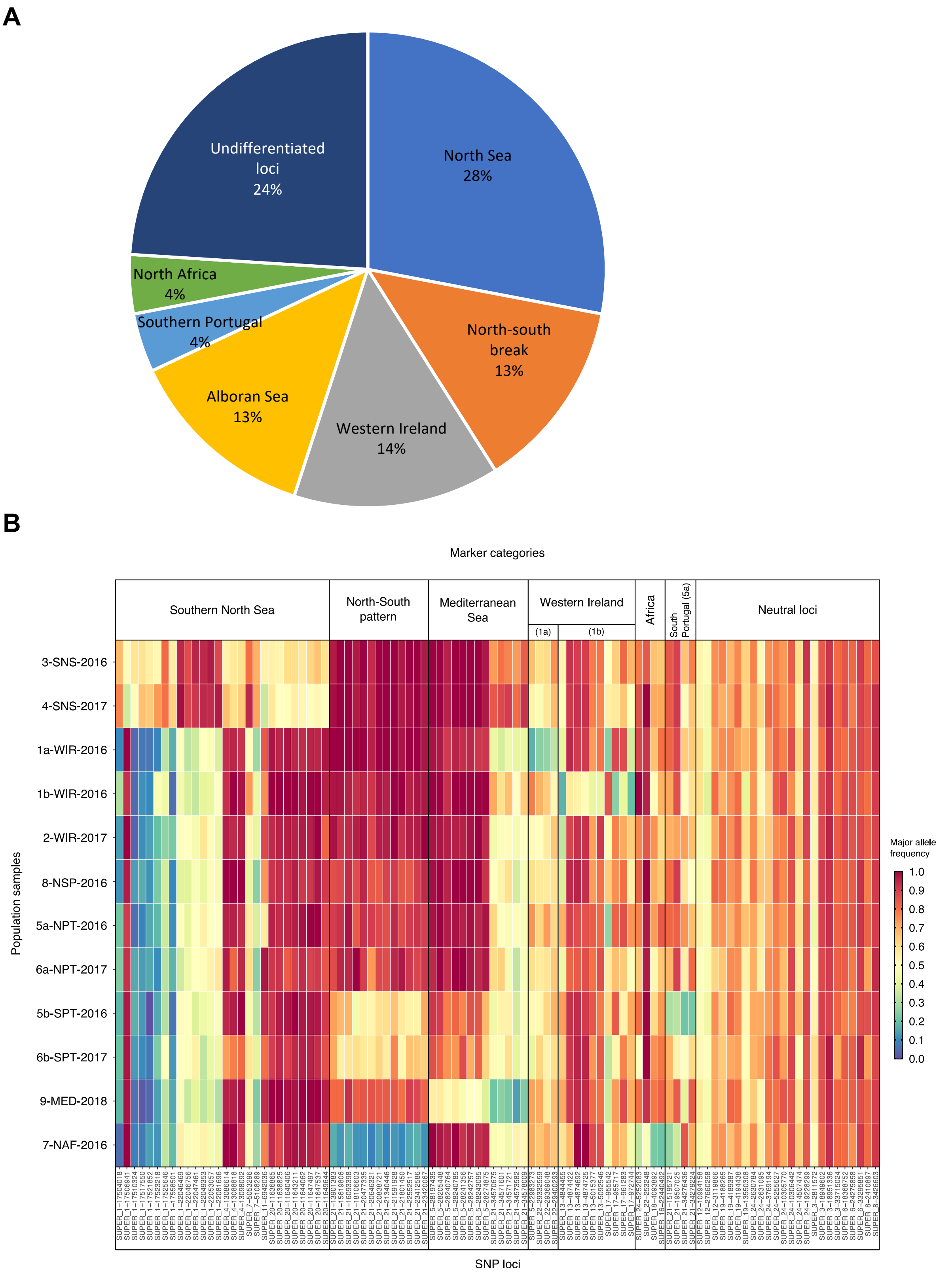


**Figure S7.** The top 100 SNPs panel. (**A**) SNP split. (**B**) Heatmap plot representing the population allele frequencies of the 100 genetic markers included in the SNP panel. Rows correspond to samples and columns to SNP loci.

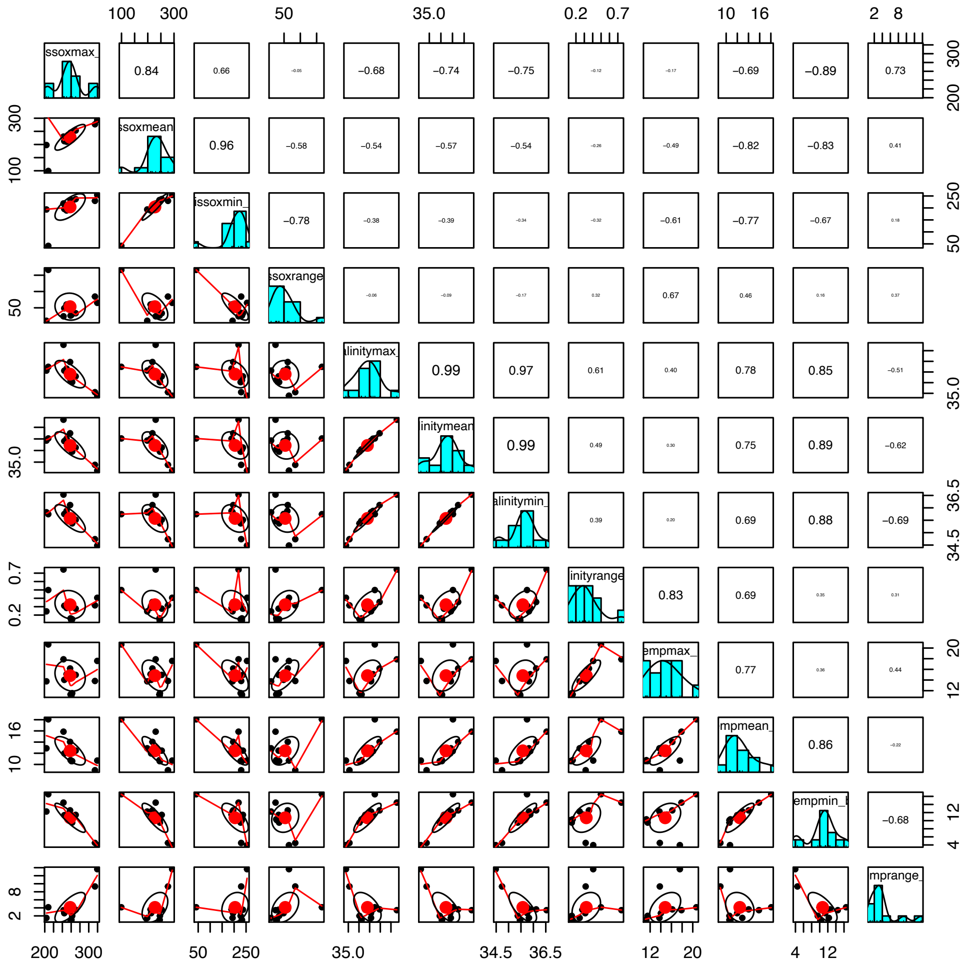

**Figure S8**. Pairwise correlation coefficients computed with the function *pairs.panels* of the R package *psych* to assess collinearity between environmental variables prior to perform a RDA.

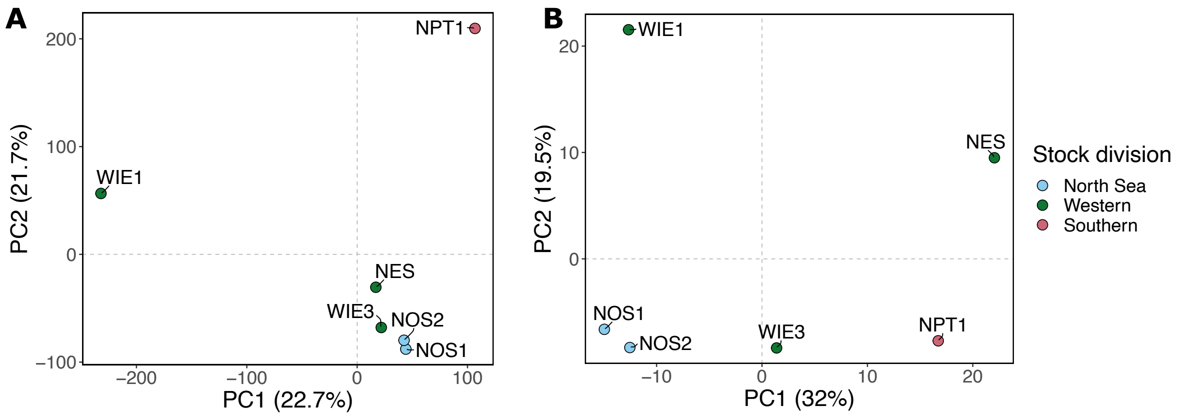


**Figure S9**. Population structure within the “northern” group. PCA plot (**A**) based on neutral markers, or (**B**) based on selective markers.

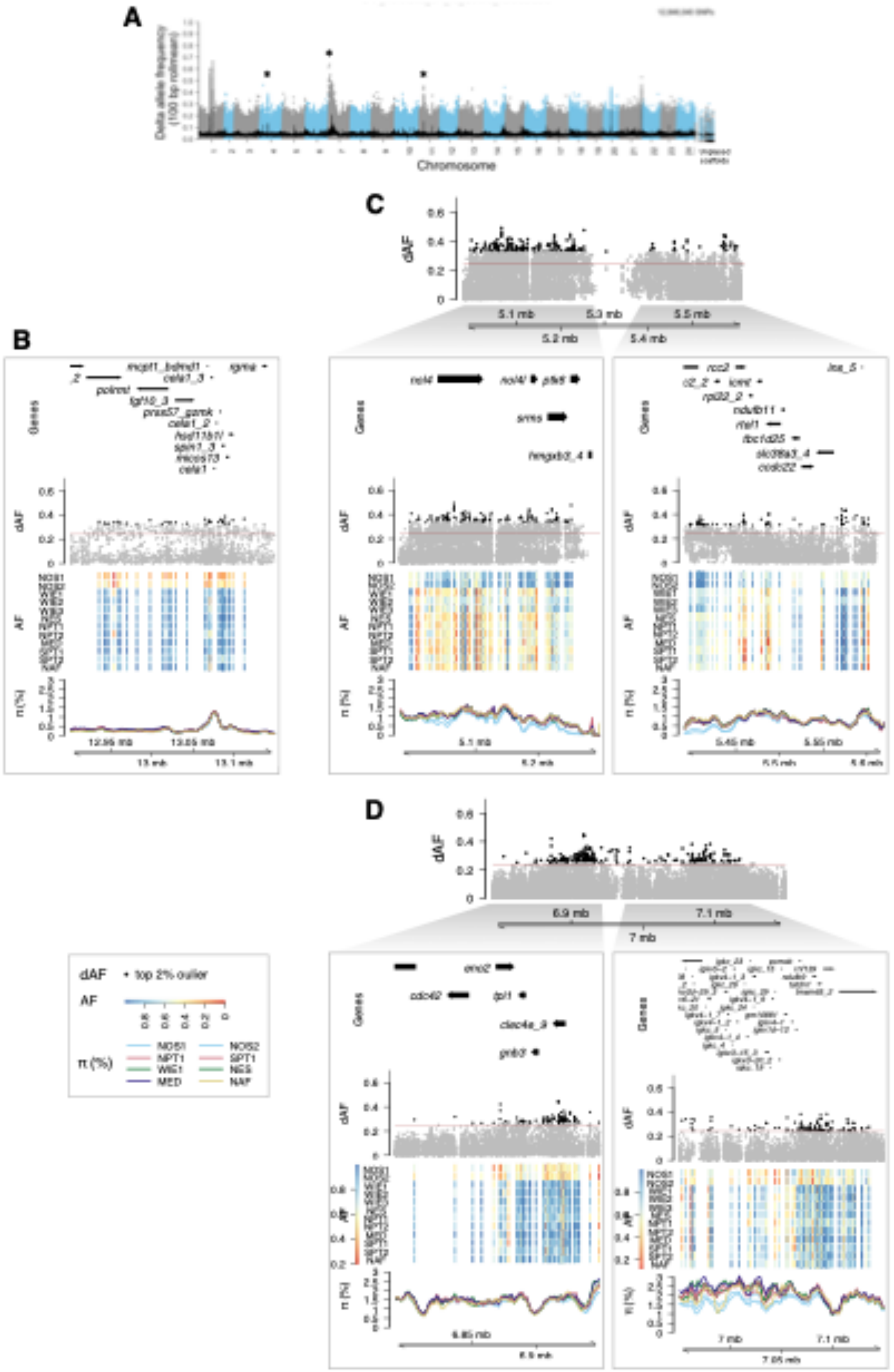


**Figure S10. Additional selective regions characteristic of the North Sea.** (**A**) Manhattan plot showing the dAF of each SNP along the genome for the contrast between the North Sea vs. all other samples. Each dot is a single SNP. The line in black is the rolling mean of dAF over 100 SNPs. Regions of interest are indicated with an arrow. Close-up plots of divergent regions in (**B**) chr 4, (**C**) chr 7, and in (**D**) chr 11. From top to bottom, the first section illustrates the gene models. The second section shows the dAF of SNPs. The top 2% SNPs are denoted in black. The third section is a heatmap plot depicting the pool minor allele frequency of the top 2% SNPs, where each row is one pool sample and each column is one variant site. The fourth section corresponds to the percentage of nucleotide diversity, $\pi$ (%), for each pool sample. The color of each line is the stock to which each sample would be designated based on the ICES stock divisions.

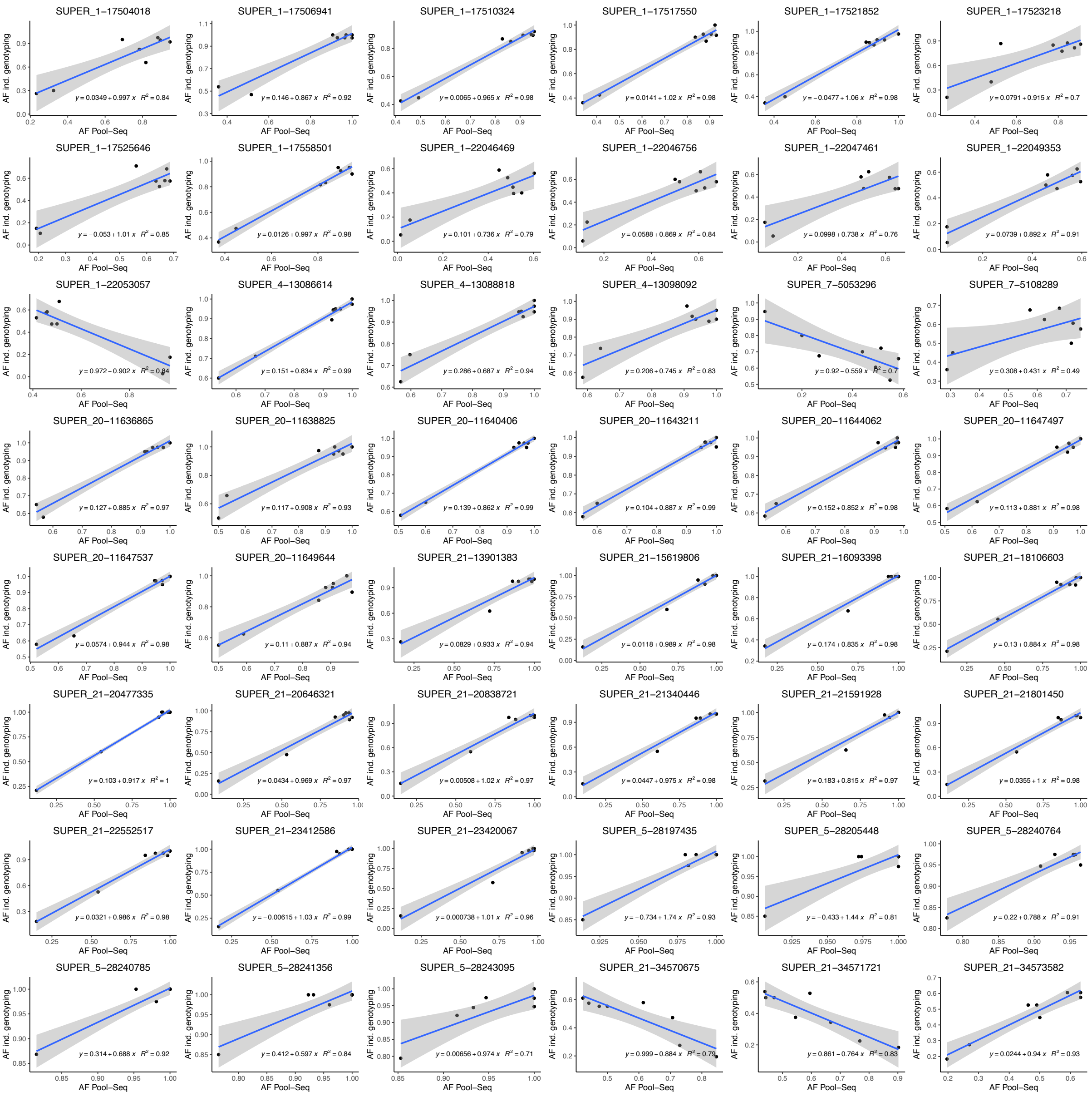


**Figure S11.** Comparison of population allele frequencies obtained with Pool-Seq and individual genotyping for the 48 SNPs putatively under selection.

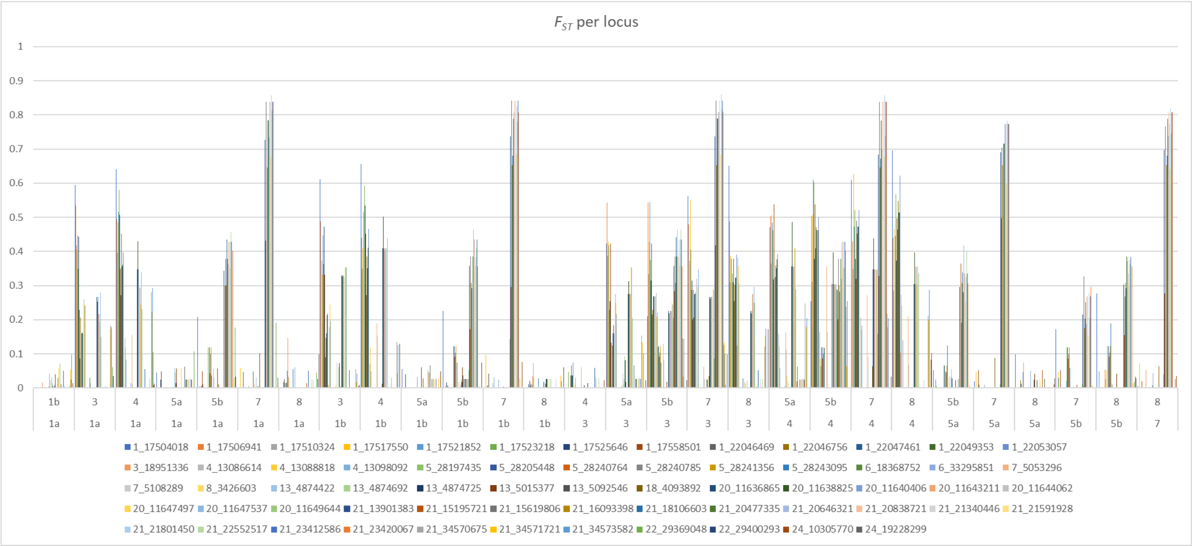


**Figure S12.** The pairwise $F_{ST}$ per locus for the 63_SNP dataset

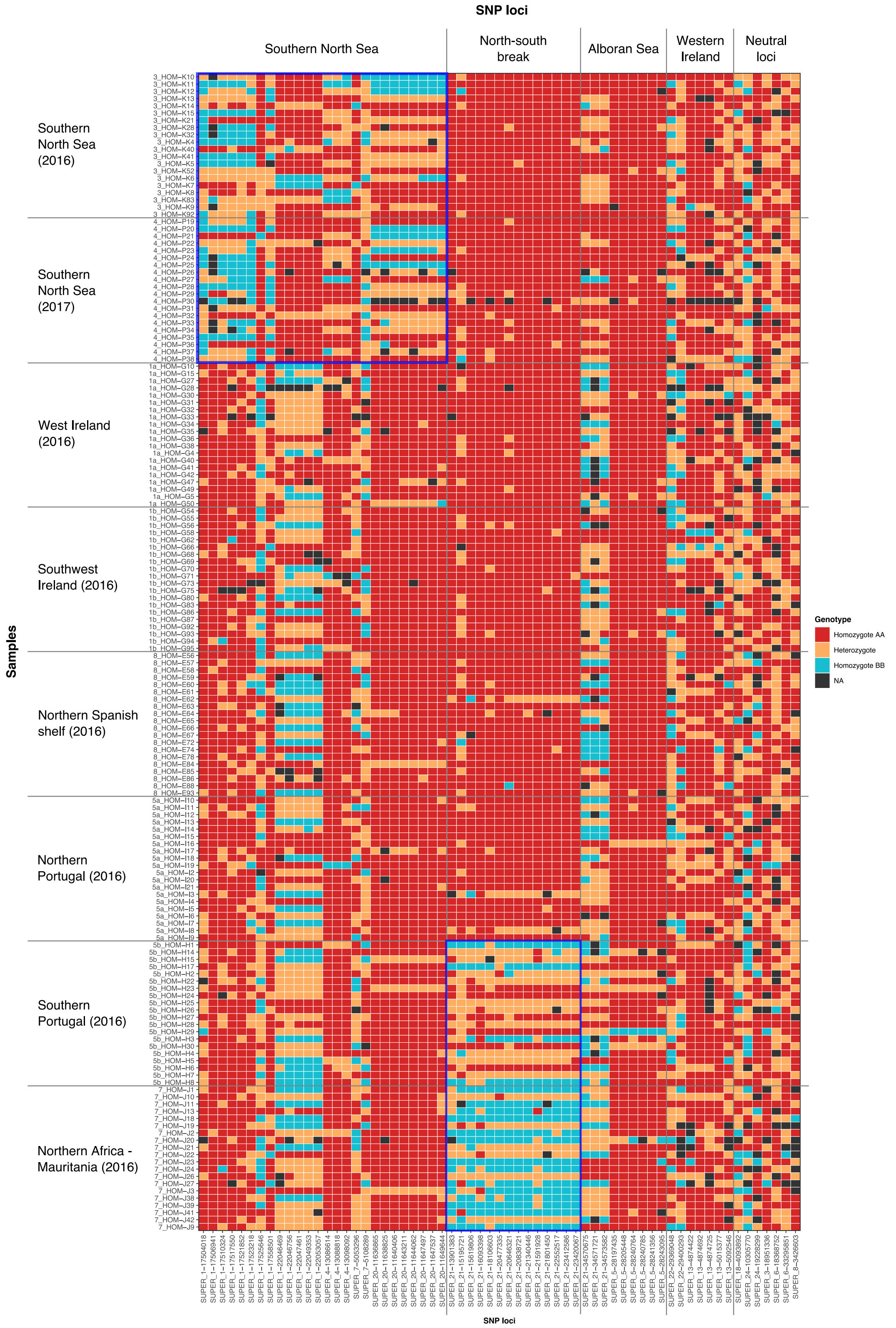


**Figure S13.** Heatmap plot representing the genotype of 157 individuals screened in 63 of the most informative SNPs for the horse mackerel. Squares in blue highlight the genotypes distinguishing the southern North Sea and the north-south genetic break.

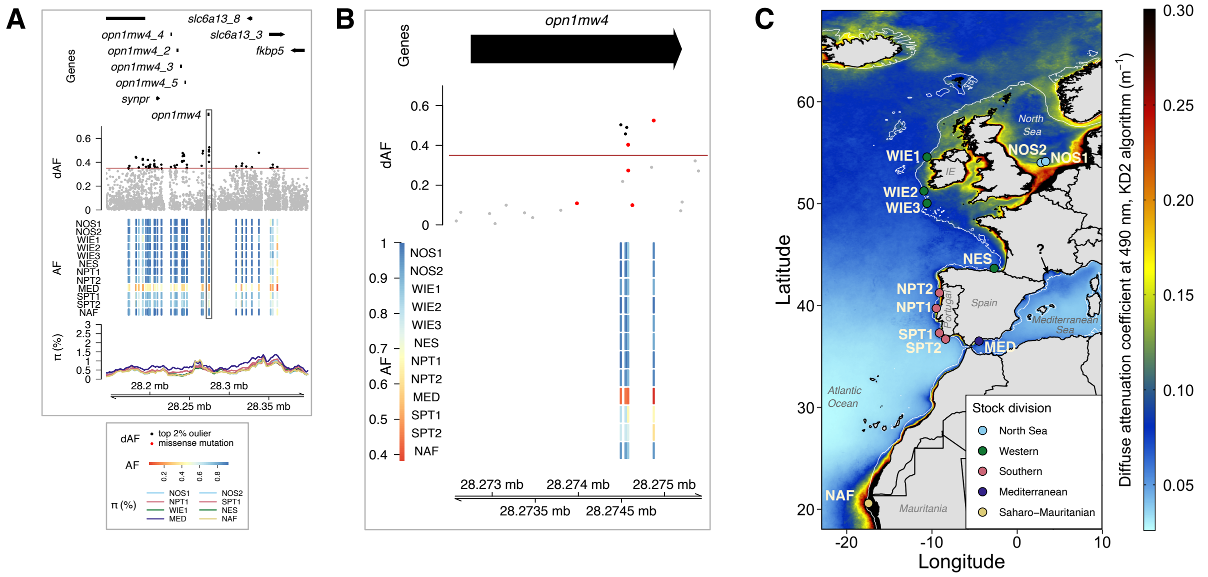

**Figure S14.** Selection signal on chr 5 that distinguishes the western Mediterranean Sea from others samples in the Atlantic Ocean. (**A**) Genomic context of the region shown in four tracks, from top to bottom: (i) gene models, (ii) delta allele frequencies (dAF) per SNP (each dot is a SNP), (iii) pool-allele frequencies for the top 2% outlier SNPs highlighted in black in the dAF track (rows are pool samples and columns are SNPs, the horizontal red line indicates the Bonferroni cutoff value of significance), and (iv) nucleotide diversity profile per pool sample. The region harboring the *opn1mw4* gene with the highest allele frequency differentiation is highlighted with a gray rectangle. (**B**) Close-up to the *opn1mw4* gene region. Missense mutations are shown as red dots in the dAF track. (**C**) Map with an overlay of the mean sea water turbidity for the east Atlantic Ocean and the western Mediterranean Sea. Turbidity data was downloaded from the OceanColor NASA MODIS-Aqua database (https://oceancolor.gsfc.nasa.gov/l3/). Turbidity values correspond to the diffuse attenuation coefficient for downwelling irradiance at 490 nm (Kd_490) in m$^{-1}$, calculated using an empirical relationship derived from *in situ* measurements of Kd_490 and blue-to-green band ratios of remote sensing reflectances (Shi & Wang, 2010). This data is a composite of mean annual values between 2002 and 2022 for a spatial resolution of 4 km. In the map, the dots denote sampling locations, and their color indicate their stock designation based on ICES 2015. Sample names as in Table 1. The arrow and '?' symbol indicate the putative spawning location of western Mediterranean populations. Isobath 200 m is shown with a white line.

**Table S1.** Collection details of the Atlantic horse mackerel samples analyzed in the current project. Abbreviations: *N*: Number of individuals, Mag: Magnetic.

| Stock | Area | Sample | Year | *N* (sample) | Latitude | Longitude | Extraction method | *N* (pool) | Pool ID | Maturity Stage | | | | | | |
|---|---|---|---|---|---|---|---|---|---|---|---|---|---|---|---|---|
| | | | | | | | | | | **1** | **2** | **3** | **4** | **5** | **6** | **NA** |
| Western | West of Ireland | 1a | 2016 | 51 | 54.42 | -10.62 | Mag Bead | 51 | WIE1 | | 31 | 19 | 1 | | | |
| Western | Southwest of Ireland | 1b | 2016 | 44 | 51.35 | -10.98 | Mag Bead | 44 | WIE2 | | 32 | 12 | | | | |
| Western | Southwest of Ireland | 2a | 2017 | 46 | 50.20 | -10.79 | Mag Bead | 62 | WIE3 | | | 44 | 2 | | | |
| Western | West of Ireland | 2b | 2017 | 16 | 53.93 | -11.09 | Mag Bead | | | | | 16 | | | | |
| North Sea | Southern North Sea | 3 | 2016 | 96 | 54.15 | 3.30 | Mag Bead | 96 | NOS1 | | 88 | | 8 | | | |
| North Sea | Southern North Sea | 4a | 2017 | 18 | 54.07 | 2.85 | Mag Bead | 70 | NOS2 | | | | 18 | | | |
| North Sea | Southern North Sea | 4b | 2017 | 21 | 54.03 | 2.90 | Mag Bead | | | | | | 21 | | | |
| North Sea | Southern North Sea | 4c | 2017 | 31 | 53.93 | 2.55 | Mag Bead | | | | | | 31 | | | |
| Southern | Northern Portugal | 5a | 2016 | 64 | 39.83 | -9.20 | Mag Bead | 64 | NPT1 | | 64 | | | | | |
| Southern | Southern Portugal | 5b | 2016 | 30 | 37.26 | -8.92 | Mag Bead | 30 | SPT1 | 22 | 5 | 3 | | | | |
| Southern | Northern Portugal | 6a | 2017 | 48 | 41.14 | -9.03 | Chelex | 47 | NPT2 | | 47 | 1 | | | | |
| Southern | Southern Portugal | 6b | 2017 | 23 | 36.84 | -8.38 | Chelex | 48 | SPT2 | | 18 | 2 | 3 | | | |
| Southern | Southern Portugal | 6c | 2017 | 25 | 36.84 | -8.10 | Chelex | | | | 19 | 6 | | | | |
| Saharo-Mauritanian | Mauritania | 7a | 2016 | 4 | 20.20 | -17.50 | Mag Bead | 57 | NAF | | 1 | | 3 | | | |
| Saharo-Mauritanian | Mauritania | 7b | 2016 | 4 | 19.00 | -17.20 | Mag Bead | | | | | | 4 | | | |
| Saharo-Mauritanian | Mauritania | 7c | 2016 | 8 | 19.90 | -17.60 | Mag Bead | | | | 1 | | 7 | | | |
| Saharo-Mauritanian | Mauritania | 7d | 2016 | 1 | 17.10 | -16.60 | Mag Bead | | | | 1 | | | | | |
| Saharo-Mauritanian | Mauritania | 7e | 2016 | 7 | 20.10 | -17.70 | Mag Bead | | | | | 1 | 6 | | | |
| Saharo-Mauritanian | Mauritania | 7f | 2016 | 4 | 20.40 | -17.70 | Mag Bead | | | | 1 | | 3 | | | |
| Saharo-Mauritanian | Mauritania | 7g | 2016 | 8 | 20.50 | -17.50 | Mag Bead | | | | 1 | | 7 | | | |
| Saharo-Mauritanian | Mauritania | 7h | 2016 | 9 | 20.50 | -17.6 | Mag Bead | | | | 4 | | 5 | | | |
| Saharo-Mauritanian | Mauritania | 7j | 2016 | 7 | 20.30 | -17.7 | Mag Bead | | | | | | 7 | | | |
| Saharo-Mauritanian | Mauritania | 7k | 2016 | 5 | 20.40 | -17.7 | Mag Bead | | | | 1 | | 4 | | | |
| Western | Northern Spanish shelf | 8a | 2016 | 22 | 43.31 | -3.46 | Mag Bead | 96 | NES | | 9 | 12 | | | | 1 |
| Western | Northern Spanish shelf | 8b | 2016 | 23 | 43.27 | -3.21 | Mag Bead | | | | 5 | 18 | | | | |
| Western | Northern Spanish shelf | 8c | 2016 | 3 | 43.27 | -2.42 | Mag Bead | | | | | 3 | | | | |
| Western | Northern Spanish shelf | 8d | 2016 | 44 | 43.22 | -2.14 | Mag Bead | | | | 15 | 28 | 1 | | | |
| Western | Northern Spanish shelf | 8e | 2016 | 4 | 43.20 | -2.10 | Mag Bead | | | | | 4 | | | | |
| Mediterranean | Alboran Sea | 9a | 2018 | 10 | 36.36 | -5.12 | CTAB | 49 | MED | | | | 10 | | | |
| Mediterranean | Alboran Sea | 9b | 2018 | 10 | 36.56 | -4.55 | CTAB | | | | | | 10 | | | |
| Mediterranean | Alboran Sea | P9c | 2018 | 10 | 36.49 | -4.42 | CTAB | | | | | | 10 | | | |
| Mediterranean | Alboran Sea | P9d | 2018 | 10 | 36.6865 | -4.28 | CTAB | | | | | | 10 | | | |
| Mediterranean | Alboran Sea | P9e | 2018 | 10 | 36.70 | -3.56 | CTAB | | | | | | 10 | | | |

**Table S2.** The international maturity scale for horse mackerel, *Trachurus trachurus.*

| Stage | Name | Female | Male |
|---|---|---|---|
| 1 | Immature | Ovaries small. Ovaries wine red and clear, torpedo shaped. | Testes small, when fresh pale flattened and transparent. When frozen it may be opaque. |
| 2 | Developing | Ovaries occupying 1/4 to almost filling body cavity. Opaque eggs visible in ovaries giving pale pink to yellow to orange coloration. Largest oocytes may have oil globules. | Gonads occupying 1/4 to to almost filing body cavity. Testes off-white to creamy white., milt not running. When frozen testes can be bleuish. |
| 3 | Spawning | Ovaries characterized by externally visible hyaline oocytes no matter how few or how early the stage of hydration. Ovary size variable from full to < 1/4 of body cavity. Ovaries can be bloodshot. | Testes from filling to < 1/4 of body cavity, milt freely running. Testes can be shrivelled (wrinkled and contracted) at anus. When frozen there might be a change of structure and the testes needs a little pushing before running. |
| 4 | Regressing Regenerating | Ovaries occupying 1/4 or less of body cavity. Ovaries reddish and often murky (dark and gloomy) in appearance, sometimes with a scattering or patch of opaque eggs. The empty ovaries will ripple when pushed together. | Ovaries occupying 1/4 or less of body cavity. Testes opaque with brownish tint and no trace of milt. When frozen testes can be bleuish ore purple. |
| 5 | Omitted spawning | No evidence of omitted spawning | No evidence of omitted spawning |
| 6 | Abnormal | No evidence of abnormal ovaries | No evidence of abnormal testes |

**Table S3.** Paired contrasts used for the calculation of delta allele frequencies. Pool IDs as in Table S4.

| Group 1 | Group 2 |
|---|---|
| Each pool | against all others |
| Southern North Sea (NOS1, NOS2) | others (WIE1, NES, NPT1, SPT1, MED) |
| West of Ireland (WIE1) | other northern samples (WIE2, NOS1, NOS2, NES, NPT1) |
| West of Ireland (WIE1, WIE2) | other northern samples (NOS1, NOS2, NES, NPT1) |
| Northern Spanish shelf (NES) | other northern samples (WIE1, NOS1, NOS2, NES, NPT1) |
| Southern Portugal and Alboran Sea (SPT1, MED) | all others (WIE1, NOS1, NOS2, NES, NPT1) |
| Southern Portugal and north Africa (SPT1, NAF) | all others (WIE1, NOS1, NOS2, NES, NPT1, MED) |
| “Northern” group (WIE1, WIE2, NOS1, NOS2, NES, NPT1) | “Southern” group (SPT1, NAF) |
| North Africa (NAF) | others (WIE1, NOS1, NOS2, NES, NPT1, SPT1, MED) |

**Table S4.** The horse mackerel samples included in the SNP validation analysis.

| Stock | Area | Sample | Pool ID | Year | # individuals | # repeated |
|---|---|---|---|---|---|---|
| Western | West of Ireland | 1a | WIE1 | 2016 | 20 | 4 |
| Western | Southwest of Ireland | 1b | WIE2 | 2016 | 20 | 4 |
| North Sea | Southern North Sea | 3 | NOS1 | 2016 | 20 | 4 |
| North Sea | Southern North Sea | 4b | NOS2 | 2017 | 20 | 4 |
| Southern | Northern Portugal | 5a | NPT1 | 2016 | 20 | 4 |
| Southern | Southern Portugal | 5b | SPT1 | 2016 | 20 | 4 |
| North African | Mauritania | 7a | NAF | 2016 | 4 | 0 |
| North African | Mauritania | 7b | NAF | 2016 | 4 | 1 |
| North African | Mauritania | 7c | NAF | 2016 | 8 | 1 |
| North African | Mauritania | 7e | NAF | 2016 | 4 | 1 |
| Western | Northern Spanish shelf | 8d | NES | 2016 | 20 | 3 |

**Table S5.** Read mapping summary statistics of the Pool-Seq data of 12 horse mackerel samples included in this study. Abbreviations: W: Western, SW: Southwestern, S: South, N: North, MQ: Mapping quality, cov.: coverage.

| Area | Pool ID | Total reads | % reads aligned | %GC | Median insert size | Mean MQ | Median cov. | Mean cov. |
|---|---|---|---|---|---|---|---|---|
| West of Ireland | WIE1 | 496,686,692 | 99.0 | 42.4 | 405 | 39.05 | 83 | 30.7 |
| Southwest of Ireland | WIE2 | 594,538,427 | 99.1 | 42.2 | 416 | 38.97 | 99 | 35.2 |
| Southwest of Ireland | WIE3 | 573,044,377 | 99.0 | 42.4 | 465 | 38.95 | 96 | 35.5 |
| Southern North Sea | NOS1 | 724,017,069 | 99.1 | 42.3 | 416 | 39 | 122 | 45.1 |
| Southern North Sea | NOS2 | 764,658,923 | 99.1 | 42.3 | 419 | 38.97 | 128 | 46.3 |
| Northern Portugal | NPT1 | 571,274,302 | 99.2 | 42.4 | 404 | 38.9 | 95 | 35.2 |
| Southern Portugal | SPT1 | 494,209,199 | 99.1 | 42.9 | 426 | 39.13 | 83 | 29.0 |
| Northern Portugal | NPT2 | 490,808,045 | 98.1 | 41.8 | *248* | 39.32 | 75 | 26.1 |
| Southern Portugal | SPT2 | 514,732,597 | 99.2 | 42.3 | *245* | 39.12 | 79 | 27.5 |
| Mauritania | NAF | 714,009,211 | 98.5 | *46.6* | 425 | 38.49 | 91 | 25.7 |
| Northern Spanish Shelf | NES | 720,020,789 | 98.9 | 43.3 | 438 | 38.96 | 122 | 41.0 |
| Alboran Sea | MED | 671,149,600 | 98.8 | 42.5 | 422 | 35.13 | 112 | 41.5 |

**Table S6.** Annotations of top 2% SNPs in divergent genomic regions. It includes distance to the closest gene (up 40 bp upstream and downstream), gene names, gene descriptions of the candidate genes, and putative type of mutation as inferred with snpEff. (This file is available as a separate Excel file due to its large size).

**Table S7.** Details of the 100 SNPs tested in the validation analyses. The SNPs highlighted in red did not reach the 80% genotyping success threshold or failed to amplify. The SNPs highlighted in orange deviated from HWE, were not polymorphic or had scoring errors and were removed from the analyses. '*LD*' indicates significant linkage disequilibrium between samples and '*Assumed*' indicates assumed LD based on chromosome position. * indicates SNPs that were included in the 17 SNP dataset.

| SNP Name | >80% success | Chr | Position | Contrast | LD Group group | Comment |
|---|---|---|---|---|---|---|
| 1_17504018* | Yes | 1 | 17504018 | Southern North Sea | Assumed | |
| 1_17506941 | Yes | 1 | 17506941 | Southern North Sea | LD | |
| 1_17510324 | Yes | 1 | 17510324 | Southern North Sea | LD | |
| 1_17517550 | Yes | 1 | 17517550 | Southern North Sea | LD | |
| 1_17521852 | Yes | 1 | 17521852 | Southern North Sea | LD | |
| 1_17523218 | Yes | 1 | 17523218 | Southern North Sea | LD | |
| 1_17525646 | Yes | 1 | 17525646 | Southern North Sea | Assumed | |
| 1_17558501 | Yes | 1 | 17558501 | Southern North Sea | LD | |
| 1_22046469 | Yes | 1 | 22046469 | Southern North Sea | LD | |
| 1_22046756 | Yes | 1 | 22046756 | Southern North Sea | LD | |
| 1_22047461 | Yes | 1 | 22047461 | Southern North Sea | LD | |
| 1_22049353 | Yes | 1 | 22049353 | Southern North Sea | LD | |
| 1_22053057* | Yes | 1 | 22053057 | Southern North Sea | LD | |
| 1_22081696 | No | 1 | 22081696 | Southern North Sea | Assumed | |
| 3_2811572 | No | 3 | 2811572 | Neutral markers | | |
| 3_18949602 | No | 3 | 18949602 | Neutral markers | | |
| 3_18951336 | Yes | 3 | 18951336 | Neutral markers | | |
| 3_33715024 | No | 3 | 33715024 | Neutral markers | | |
| 4_13086614* | Yes | 4 | 13086614 | Southern North Sea | LD | |
| 4_13088818 | Yes | 4 | 13088818 | Southern North Sea | LD | |
| 4_13098092 | Yes | 4 | 13098092 | Southern North Sea | LD | |
| 5_22983273 | No | 5 | 22983273 | Western Ireland (1a) | | |
| 5_28197435 | Yes | 5 | 28197435 | Med and/or S Portugal | | |
| 5_28205448 | Yes | 5 | 28205448 | Med and/or S Portugal | | |
| 5_28240764 | Yes | 5 | 28240764 | Med and/or S Portugal | | |
| 5_28240785 | Yes | 5 | 28240785 | Med and/or S Portugal | | |
| 5_28241356* | Yes | 5 | 28241356 | Med and/or S Portugal | | |
| 5_28242757 | No | 5 | 28242757 | Med and/or S Portugal | | |
| 5_28243095 | Yes | 5 | 28243095 | Med and/or S Portugal | | |
| 5_28274875 | No | 5 | 28274875 | Med and/or S Portugal | | |
| 6_18368752* | Yes | 6 | 18368752 | Neutral markers | | |
| 6_24275858 | No | 6 | 24275858 | Neutral markers | | |
| 6_33295851* | Yes | 6 | 33295851 | Neutral markers | | |
| 7_5053296* | Yes | 7 | 5053296 | Southern North Sea | | |
| 7_5108289 | Yes | 7 | 5108289 | Southern North Sea | | |
| 8_2410897 | No | 8 | 2410897 | Neutral markers | | |
| 8_3426603* | Yes | 8 | 3426603 | Neutral markers | | |
| 11_6942036 | Yes | 11 | 6942036 | Southern North Sea | | Out of HWE in 2 pops |
| 12_3119866 | Yes | 12 | 3119866 | Neutral markers | | Not polymorphic |
| 12_10994158 | No | 12 | 10994158 | Neutral markers | | |
| 12_27660258 | Yes | 12 | 27660258 | Neutral markers | | Out of HWE in 3 pops |
| 13_4844455 | No | 13 | 4844455 | Western Ireland (1b) | | |
| 13_4874422 | Yes | 13 | 4874422 | Western Ireland (1b) | LD | |
| 13_4874692 | Yes | 13 | 4874692 | Western Ireland (1b) | LD | |
| 13_4874725 | Yes | 13 | 4874725 | Western Ireland (1b) | LD | |
| 13_5015377* | Yes | 13 | 5015377 | Western Ireland (1b) | | |
| 13_5092546 | Yes | 13 | 5092546 | Western Ireland (1b) | | |
| 16_22440492 | No | 16 | 22440492 | Africa | | |
| 17_955542 | No | 17 | 955542 | Western Ireland (1b) | | |
| 17_955717 | Yes | 17 | 955717 | Western Ireland (1b) | | Out of HWE in 1 pop |
| 17_961283 | No | 17 | 961283 | Western Ireland (1b) | | |

**Table S7.** (*Continuation*).

| SNP Name | >80% success | Chr | Position | Contrast | LD Group group | Comment |
|---|---|---|---|---|---|---|
| 17_972744 | Yes | 17 | 972744 | Western Ireland (1b) | | Not polymorphic |
| 18_4093892* | Yes | 18 | 4093892 | Africa | | |
| 19_4188265 | No | 19 | 4188265 | Neutral markers | | |
| 19_4189387 | No | 19 | 4189387 | Neutral markers | | |
| 19_4194438 | No | 19 | 4194438 | Neutral markers | | |
| 19_13550308 | No | 19 | 13550308 | Neutral markers | | |
| 20_11636865 | Yes | 20 | 11636865 | Southern North Sea | LD | |
| 20_11638825* | Yes | 20 | 11638825 | Southern North Sea | LD | |
| 20_11640406 | Yes | 20 | 11640406 | Southern North Sea | LD | |
| 20_11643211 | Yes | 20 | 11643211 | Southern North Sea | LD | |
| 20_11644062 | Yes | 20 | 11644062 | Southern North Sea | LD | |
| 20_11647497 | Yes | 20 | 11647497 | Southern North Sea | LD | |
| 20_11647537 | Yes | 20 | 11647537 | Southern North Sea | LD | |
| 20_11649644 | Yes | 20 | 11649644 | Southern North Sea | LD | |
| 21_13901383 | Yes | 21 | 13901383 | North-South pattern | | |
| 21_15195721 | Yes | 21 | 15195721 | Southern Portugal | | |
| 21_15619806* | Yes | 21 | 15619806 | North-South pattern | | |
| 21_16093398 | Yes | 21 | 16093398 | North-South pattern | | |
| 21_18106603 | Yes | 21 | 18106603 | North-South pattern | | |
| 21_19507025 | Yes | 21 | 19507025 | Southern Portugal | | Out of HWE in 1 pop |
| 21_20477335 | Yes | 21 | 20477335 | North-South pattern | | |
| 21_20646321 | Yes | 21 | 20646321 | North-South pattern | LD | |
| 21_20838721 | Yes | 21 | 20838721 | North-South pattern | LD | |
| 21_21340446 | Yes | 21 | 21340446 | North-South pattern | LD | |
| 21_21591928 | Yes | 21 | 21591928 | North-South pattern | | |
| 21_21801450 | Yes | 21 | 21801450 | North-South pattern | | |
| 21_22552517 | Yes | 21 | 22552517 | North-South pattern | | |
| 21_23412586* | Yes | 21 | 23412586 | North-South pattern | LD | |
| 21_23420067 | Yes | 21 | 23420067 | North-South pattern | LD | |
| 21_34276436 | No | 21 | 34276436 | Southern Portugal | | |
| 21_34279224 | No | 21 | 34279224 | Southern Portugal | | |
| 21_34570675 | Yes | 21 | 34570675 | Med and/or S Portugal | LD | |
| 21_34571601 | No | 21 | 34571601 | Med and/or S Portugal | | |
| 21_34571721 | Yes | 21 | 34571721 | Med and/or S Portugal | LD | |
| 21_34573582* | Yes | 21 | 34573582 | Med and/or S Portugal | LD | |
| 21_34578009 | No | 21 | 34578009 | Med and/or S Portugal | | |
| 22_253248 | No | 22 | 253248 | Africa | | |
| 22_29332559 | Yes | 22 | 29332559 | Western Ireland (1a) | | Out of HWE in 5 pops |
| 22_29369048* | Yes | 22 | 29369048 | Western Ireland (1a) | | |
| 22_29400293 | Yes | 22 | 29400293 | Western Ireland (1a) | | |
| 24_2630784 | No | 24 | 2630784 | Neutral markers | | |
| 24_2631095 | No | 24 | 2631095 | Neutral markers | | |
| 24_3769194 | No | 24 | 3769194 | Neutral markers | | |
| 24_5252083 | Yes | 24 | 5252083 | Africa | | Scoring error |
| 24_5255627 | No | 24 | 5255627 | Neutral markers | | |
| 24_10305770* | Yes | 24 | 10305770 | Neutral markers | | |
| 24_10306442 | Yes | 24 | 10306442 | Neutral markers | | Out of HWE in 1 pop |
| 24_14507474 | No | 24 | 14507474 | Neutral markers | | |
| 24_19228299* | Yes | 24 | 19228299 | Neutral markers | | |

**Table S8.** Pairwise multi-locus $F_{ST}$ (above the diagonal) and associated *P*-values (below the diagonal) for the 63_SNP dataset (top panel) 17_SNP dataset (bottom panel). *P*-values highlighted in red were still significant after sequential Bonferroni correction.

| | 1a | 1b | 3 | 4 | 5a | 5b | 7 | 8 |
|---|---|---|---|---|---|---|---|---|
| 1a | | 0.004 | 0.195 | 0.260 | -0.004 | 0.135 | 0.361 | 0.004 |
| 1b | 0.28 | | 0.198 | 0.265 | -0.006 | 0.124 | 0.352 | -0.003 |
| 3 | 0.00 | 0.00 | | -0.006 | 0.180 | 0.243 | 0.417 | 0.218 |
| 4 | 0.00 | 0.00 | 0.60 | | 0.241 | 0.287 | 0.446 | 0.286 |
| 5a | 0.61 | 0.71 | 0.00 | 0.00 | | 0.101 | 0.323 | -0.002 |
| 5b | 0.00 | 0.00 | 0.00 | 0.00 | 0.00 | | 0.080 | 0.111 |
| 7 | 0.00 | 0.00 | 0.00 | 0.00 | 0.00 | 0.00 | | 0.334 |
| 8 | 0.29 | 0.54 | 0.00 | 0.00 | 0.51 | 0.00 | 0.00 | |

| | 1a | 1b | 3 | 4 | 5a | 5b | 7 | 8 |
|---|---|---|---|---|---|---|---|---|
| 1a | | 0.016 | 0.138 | 0.196 | 0.004 | 0.088 | 0.241 | 0.009 |
| 1b | 0.09 | | 0.121 | 0.190 | 0.003 | 0.075 | 0.221 | -0.002 |
| 3 | 0.00 | 0.00 | | -0.002 | 0.102 | 0.137 | 0.297 | 0.168 |
| 4 | 0.00 | 0.00 | 0.53 | | 0.154 | 0.187 | 0.340 | 0.233 |
| 5a | 0.32 | 0.35 | 0.00 | 0.00 | | 0.033 | 0.183 | 0.006 |
| 5b | 0.00 | 0.00 | 0.00 | 0.00 | 0.01 | | 0.055 | 0.068 |
| 7 | 0.00 | 0.00 | 0.00 | 0.00 | 0.00 | 0.00 | | 0.209 |
| 8 | 0.17 | 0.52 | 0.00 | 0.00 | 0.26 | 0.00 | 0.00 | |
